# HNF1α transcriptional activation and repression maintain human islet α and β cell function

**DOI:** 10.1101/2022.09.25.509394

**Authors:** Mollie F. Qian, Romina J. Bevacqua, Vy M. Nguyen, Xiong Liu, Weichen Zhao, Charles A. Chang, Xueying Gu, Xiao-Qing Dai, Patrick E. MacDonald, Seung K. Kim

## Abstract

*HNF1A* haploinsufficiency underlies the most common form of human monogenic diabetes (HNF1A-MODY) and hypomorphic *HNF1A* variants confer type 2 diabetes risk, but a lack of experimental systems has limited our understanding of how the transcription factor HNF1α regulates adult human islet function. Here, we combined human islet genetics, RNA sequencing, Cleavage Under Targets & Release Using Nuclease (CUT&RUN) chromatin mapping, patch-clamp electrophysiology and transplantation-based assays to elucidate HNF1α-regulated mechanisms in mature pancreatic α and β cells. shRNA-mediated suppression of *HNF1A* in primary human pseudoislets led to blunted insulin output and dysregulated glucagon secretion both *in vitro* and after transplantation into immunocompromised mice, recapitulating phenotypes observed in HNF1A-MODY patients. These deficits corresponded with altered expression of genes encoding factors critical for hormone secretion, including calcium channel subunits, ATP-transporters and extracellular matrix constituents. Additionally, *HNF1A* loss led to upregulation of transcriptional repressors, providing evidence for a mechanism of transcriptional de-repression through HNF1α. CUT&RUN mapping of HNF1α DNA-binding sites in primary human islets verified that a subset of HNF1α-regulated genes were direct targets. These data provide unprecedented mechanistic links between *HNF1A* loss and diabetic phenotypes in mature human α and β cells.

## INTRODUCTION

Human diabetes genetics has been advanced by studies of monogenic diabetes, like Maturity Onset Diabetes of the Young (MODY) (Zhang et al. 2021), and by identification of causal variants of type 2 diabetes (T2D) through genome-wide association studies (GWAS) (Mahajan et al. 2018). The most common form of MODY (HNF1A-MODY) results from mutations in *HNF1A*, which encodes the transcription factor HNF1α (Riddle et al. 2020), and multiple hypomorphic *HNF1A* variants have been associated with increased T2D risk (Najmi et al. 2017; Althari et al. 2020). Despite the strong association between *HNF1A* deficiency and human diabetes, the mechanisms by which HNF1α regulates mature human islet cell function remain poorly understood.

Previous studies of HNF1A-MODY patients have revealed impaired insulin secretion that improves with sulfonylurea treatment (Bacon et al. 2016; Østoft et al. 2015). These and other studies in human stem cell models (Cardenas-Diaz et al. 2019; Low et al. 2021; Cujba et al. 2022; González et al. 2022) strongly suggest a role for HNF1α in pancreatic β cell development and function. In addition to β cells, *HNF1A* is abundantly expressed in α cells (Arda et al. 2016; Blodgett et al. 2015), consistent with observations that HNF1A-MODY patients have dysregulated glucagon secretion (Haliyur et al. 2019; Tuomi et al. 2006; Østoft et al. 2014). Evidence also supports the view that α cell dysfunction governs disease development and severity in type 1 diabetes (T1D) and T2D (Gromada et al. 2018), but the impact of *HNF1A* loss in mature human α cells is unknown.

Deciphering HNF1α function in adult islet cells has been complicated by challenges of assessing *Hnf1a* function in mouse models. In humans, *HNF1A* haploinsufficiency underlies HNF1A-MODY, but mice heterozygous for an *Hnf1a* null allele are not diabetic (Pontoglio et al. 1998). Studies of homozygous germline *Hnf1a* null mice are confounded by severe hepatic and renal phenotypes not observed in humans (Pontoglio et al. 1998). These limitations, and the realization that *HNF1A* deficiency leads to functional deficits in human β and α cells (Haliyur et al. 2019), motivated development of human islet systems to study HNF1α function. Studies using *HNF1A* deficient β-like cells derived from human multipotent stem cells have characterized HNF1α regulation during islet cell development (Cardenas-Diaz et al. 2019; Low et al. 2021; Cujba et al. 2022; González et al. 2022), but progeny cells from these studies did not fully recapitulate adult β cell functions, limiting conclusions about the roles of HNF1α in mature adult islets. Functional and transcriptional characterization of islets from an HNF1A-MODY patient were recently reported (Haliyur et al. 2019), but conclusions in this case were inferred from studies of a single subject.

To elucidate mechanisms by which HNF1α loss leads to impaired human pancreatic α and β cell function, we combined modern human islet genetics, RNA sequencing, Cleavage Under Targets & Release Using Nuclease (CUT&RUN) genomic mapping, patch-clamp electrophysiology, and transplantation-based assays (Bevacqua et al. 2021b, 2021a; Skene and Henikoff 2017; Tellez et al. 2020; Lin et al. 2021; Enge et al. 2017). These approaches applied the advantages of genetic and physiological studies in human ‘pseudoislets’, formed by sequential dispersion and re-aggregation of primary human islets to permit efficient genetic targeting (reviewed in Friedlander et al. 2021). Together, our findings provide evidence that HNF1α directly regulates genes essential for regulated hormone secretion in human adult α and β cells.

## RESULTS

### Loss of *HNF1A* in primary human islets leads to impaired islet cell function

To achieve conditional *HNF1A* loss in primary human islets, we expressed shRNA targeting *HNF1A* with a lentiviral vector that co-expressed a GFP reporter (Fig. 1A: Materials and Methods). Lentivirus expressing shRNA that suppressed *HNF1A* (HNF1AKD; ‘knockdown’, KD) or non-targeting control shRNA were delivered to dispersed adult islet cells (Supplemental Table S1), which were re-aggregated to form pseudoislets (Fig. 1B: Materials and Methods). At 5 days post-lentiviral infection, this approach yielded significant, reproducible reduction in *HNF1A* mRNA assessed by RT-qPCR (Fig. 1C). We also observed reduced expression of the putative HNF1α targets *HNF4A, HNF1A-AS1, TMEM27, KCNJ11*, and *SLC2A2* (Fig. 1D).

**Figure 1.**
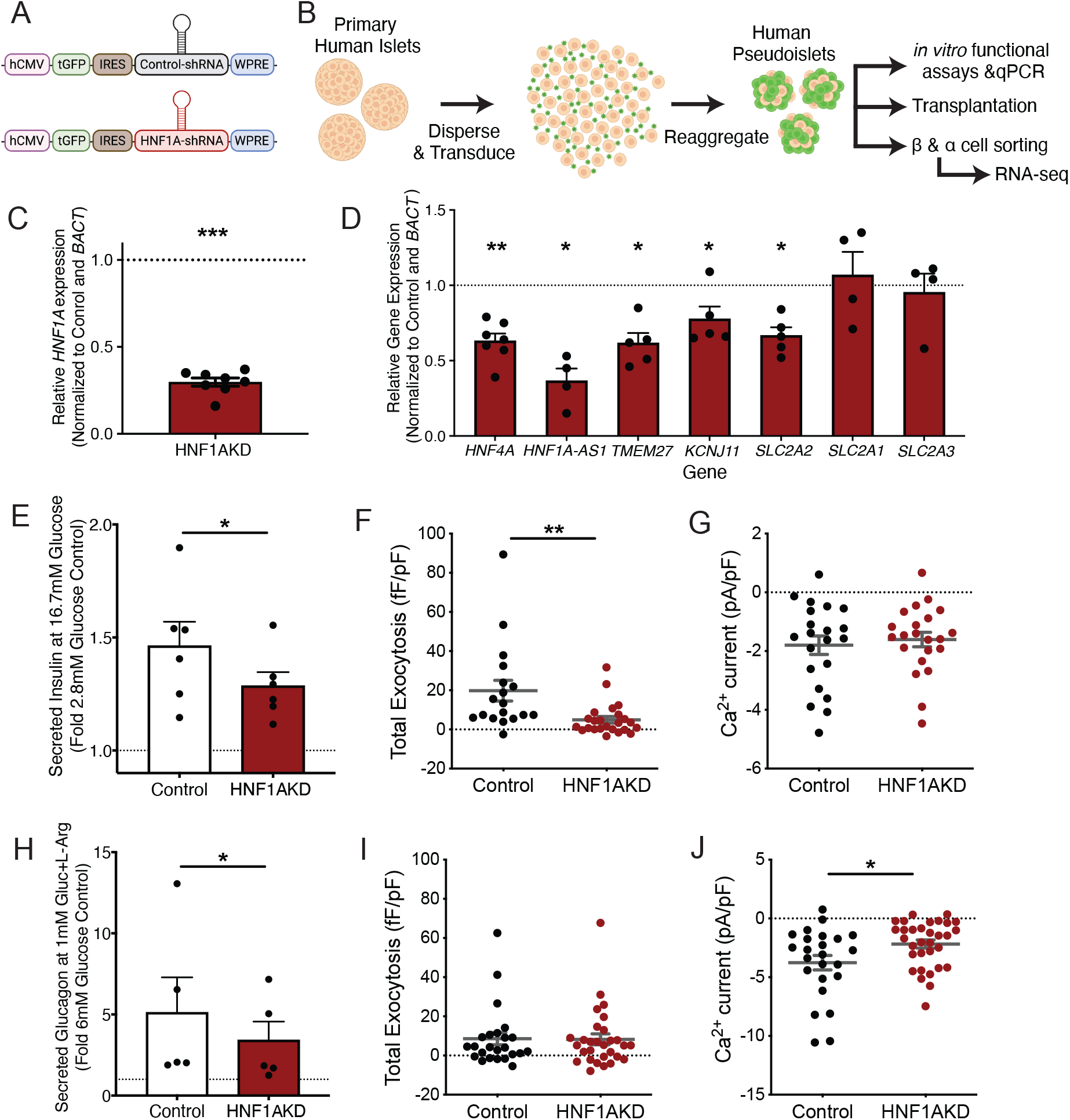
shRNA-mediated ‘knockdown’ (KD) of *HNF1A* in primary human islets results in acute dysregulation of islet cell function. (A) Schematic of lentiviral constructs coding for shRNA and a GFP reporter (tGFP); Control-shRNA= non-targeting (“Control”), HNF1A-shRNA= HNF1A-targeting (“HNF1AKD”). (B) Formation of pseudoislets for downstream assays; transduction with lentivirus, followed by reaggregation over 5 days in culture. (C, D) mRNA expression of (C) *HNF1A* and (D) putative HNF1α targets in HNF1AKD relative to Control pseudoislets; statistics performed on deltaCT values (n=4-8 donors per gene). Secreted (E) insulin (n=6 donors) and (H) glucagon (n=5 donors) from pseudoislet static batch assays; Gluc=Glucose and L-Arg=10mM L-Arginine. (F-G, I-J) Patch-clamp electrophysiology recordings at 5mM glucose of (F, I) total exocytosis and (G, J) early calcium currents from (F-G) β and (I-J) α cells (n=18-32 cells per condition from 3 human donors). Data are presented as mean values ± SEM. Two-tailed t-tests were used to generate *P*-values; \**P*<0.05, \*\**P*<0.01, \*\*\**P*<0.00001.

We next assessed hormone secretion after acute *HNF1A* loss in primary islets. In static batch assays, insulin secretion from HNF1AKD pseudoislets was significantly reduced at 16.7 mM glucose (Fig. 1E). In agreement with these findings, patch-clamp electrophysiology (Fig. 1F-G, I-J) revealed reduced exocytosis in HNF1AKD β cells (Fig. 1F, Supplemental Fig. S1A). Likewise, glucagon secretion from HNF1AKD pseudoislets was blunted after stimulation with 1mM glucose+10mM L-Arginine (Fig. 1H). Moreover, patch-clamp of HFN1AKD α cells revealed reduced calcium (Ca^2+^) currents (Fig. 1J, Supplemental Fig. S1B), although at the glucose concentration used here exocytosis was already suppressed in these cells as we recently showed (Dai et al. 2022). While insulin transcript levels were reduced, insulin and glucagon protein expression were not significantly different in HNF1AKD versus control pseudoislets (Supplemental Fig. S1C-F). These physiological studies provide evidence of dysregulated human α and β cell function after acute *HNF1A* loss in adult primary islets.

### HNF1α deficiency leads to *in vivo* α and β cell phenotypes observed in HNF1A-MODY

To investigate the durability of physiological defects observed in pseudoislets after suppression of *HNF1A*, we transplanted control and HNF1AKD pseudoislets into the renal capsules of appropriate mouse models. To study defects of insulin output, we transplanted pseudoislets into immunocompromised, nondiabetic *NOD scid IL2Rγ*^*null*^ (NSG) mice, a strain permitting xenotransplantation of tissues like islets (Materials and Methods). One month after transplantation, we measured human β cell function using a human-specific insulin ELISA (Fig. 2A). After intraperitoneal glucose tolerance testing (IPGTT), we observed significant blunting of insulin secretion from HNF1AKD grafts (Fig. 2B-D). This deficit was ameliorated by treatment of mice with glibenclamide (Supplemental Fig. S2A-D), a sulfonylurea used to treat insulin insufficiency in HNF1A-MODY patients. After recovery of human pseudoislet grafts, immunostaining demonstrated that HNF1α expression was significantly reduced in HNF1AKD grafts (Supplemental Fig. S2E-G), confirming that *HNF1A* suppression was sustained after months *in vivo*. Thus, acute *HNF1A* deficiency in human adult pseudoislets was durable, and phenocopied key elements of insulin insufficiency observed in *HNF1A* deficient patients.

**Figure 2.**
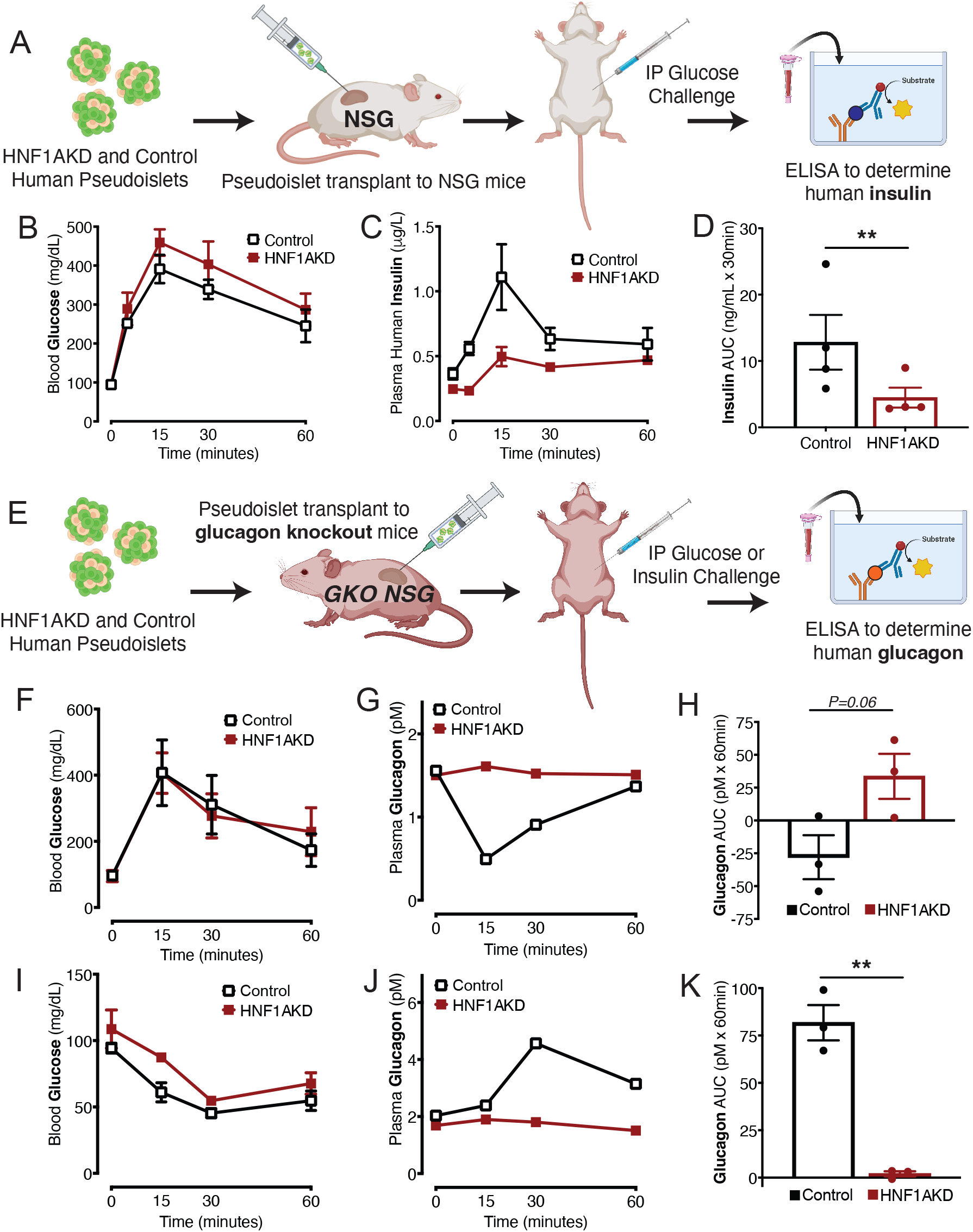
*HNF1A* suppression leads to dysregulated insulin and glucagon secretion after one month *in vivo*. (A) Experimental approach for control and HNF1AKD pseudoislet transplantation to kidney capsules of NSG mice and characterization of *in vivo* phenotypes. (B) Blood glucose, (C) plasma human insulin levels, and (D) area under the curve (AUC) of insulin excursion upon intraperitoneal (IP) glucose challenge after transplantation of pseudoislets to NSG mice (n=4 mice, 3 human donors). (E) Schematic of pseudoislet transplantation to glucagon knockout mice on an NSG background (GKO NSG) for characterization of *in vivo* glucagon phenotypes. (F, I) Blood glucose, (G, J) representative plasma glucagon levels, and (H, K) AUC of glucagon excursion upon IP (F-H) glucose or (I-K) insulin challenge after transplantation to GKO NSG mice (n=3 mice, 3 human donors). Data are mean values ± SEM. Two-tailed t-tests were used to generate *P*-values; \*\**P*<0.01.

In addition to impaired insulin output, dysregulated glucagon secretion has also been documented in HNF1A-MODY patients and in primary islets isolated from an HNF1A-MODY donor (Østoft et al. 2014; Haliyur et al. 2019). Due to complete conservation of glucagon between mouse and human, it is not possible to distinguish between human graft-derived and endogenous mouse host glucagon after transplantation of pseudoislets in the NSG strain. To overcome this barrier, we used a glucagon knockout (GKO) NSG mouse model (hereafter, GKO NSG) (Tellez et al. 2020) that harbors an in-frame deletion of *Glucagon1-29*, eliminating endogenous glucagon. This permits ELISA-based detection of glucagon produced by transplanted human α cells (Fig. 2E). We transplanted control and HNF1AKD pseudoislets in GKO NSG mice, then assessed glucagon secretion through IPGTT one month after transplantation. Glucose challenge resulted in comparable levels of acute hyperglycemia in GKO NSG mice transplanted with control or HNF1AKD pseudoislets (Fig. 2F). As expected, glucagon secretion from control grafts decreased in response to hyperglycemia by 30 minutes after glucose injection (Fig. 2G, Supplemental Fig. S3A-C). In contrast, glucagon secretion was not suppressed in HNF1AKD grafts after glucose challenge, and area under the curve (AUC) measures of net glucagon secretion were higher for HNF1AKD versus control grafts (Fig. 2H). These findings are reminiscent of the inappropriate increase in glucagon secretion observed after glucose challenge in HNF1A-MODY patients (Tuomi et al. 2006; Østoft et al. 2014).

In addition to glucagon hypersecretion at high glucose levels, studies of islets from an HNF1A-MODY donor documented reduced glucagon secretion in conditions that normally stimulate glucagon output (Haliyur et al. 2019). To investigate effects of *HNF1A* loss on glucagon secretion in response to hypoglycemia, we transplanted HNF1AKD pseudoislets in GKO NSG mice and performed IP insulin tolerance tests (IP-ITT). IP-ITT led to acute hypoglycemia in GKO NSG mice independent of control or HNF1AKD graft type (Fig. 2I), but glucagon output was blunted in HNF1AKD grafts (Fig. 2J-K, Supplemental Fig. S3D-F). In summary, our transplantation studies reveal that acute loss of *HNF1A* in adult primary islets phenocopied multiple hormone secretion defects observed in human HNF1A-MODY, including reduced insulin secretion, inappropriately high glucagon output during hyperglycemia, and impaired glucagon secretion during hypoglycemia.

### RNA-seq identifies transcriptome changes after *HNF1A* loss in mature β cells

HNF1α is a known transcriptional regulator (Miyachi et al. 2022); thus, to investigate the mechanisms underlying islet phenotypes detailed above, we performed RNA sequencing (RNA-seq) to characterize the β cell transcriptome after *HNF1A* loss (Fig. 3A). To isolate HNF1AKD β cells, we used fluorescence-activated cell sorting (FACS) to purify lentivirus-infected GFP^+^ cells expressing NTPDase3, a human β cell surface marker (Saunders et al. 2019). Enrichment of insulin (*INS*) mRNA in NTPDase3^+^ fractions was confirmed by RT-qPCR (Supplemental Fig. S4A-B). RNA-seq libraries of GFP^+^ NTPDase3^+^ cells from 4 human donors were produced and sequenced. Principal Component Analysis (PCA) showed clustering of samples by donor, consistent with prior studies (Arda et al. 2016; Peiris et al. 2018; Bevacqua et al. 2021b), in addition to separation of HNF1AKD and control β cells (Supplemental Fig. S4C-D). The DESeq2 algorithm (Love et al. 2014) identified 1,605 differentially expressed genes in β cells after *HNF1A* loss, corresponding to 800 genes with significantly reduced mRNA, including *HNF1A* (Fig. 3B), and 805 with increased mRNA levels (Fig. 3C-D).

**Figure 3.**
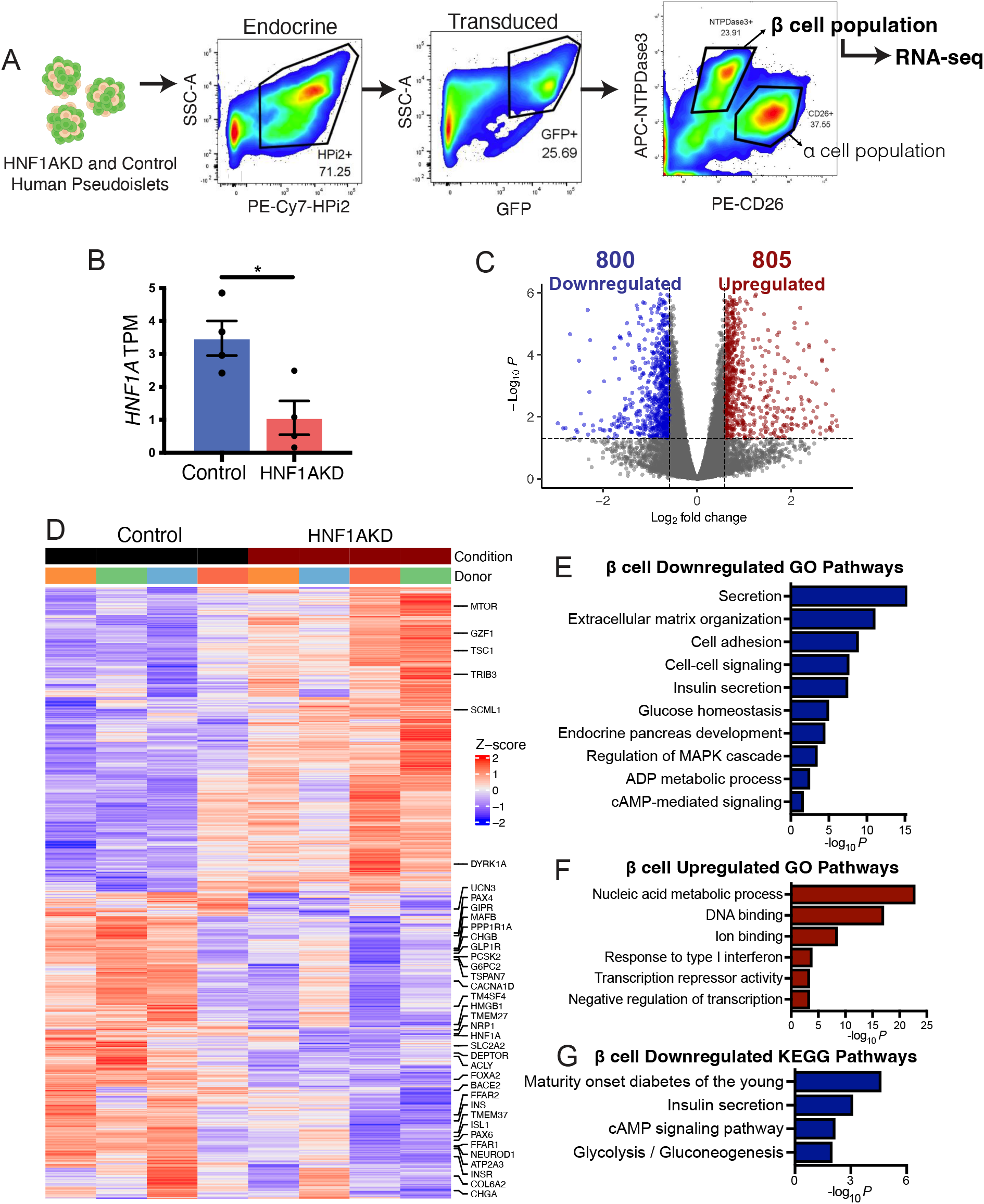
RNA-seq of HNF1AKD β cells shows that HNF1α regulates insulin secretion, metabolism, developmental pathways, and cell-cell signaling in β cells. (A) Schematic of FACS scheme for isolation of transduced live β cells (HPi2+GFP+NTPDase3+) from control and HNF1AKD pseudoislets for downstream RNA sequencing. (B) *HNF1A* transcripts per million (TPM) in sequenced samples. (C) Differential expression (DE) analysis revealed significantly up- and down-regulated genes after HNF1AKD in β cells; thresholds: Fold change (FC)= 1.5, adjusted *P*-value= 0.05. (D) Heatmap of DEGs in β cells after HNF1AKD. Significantly (E) downregulated and (F) upregulated gene ontology (GO) pathways and (G) downregulated KEGG pathways in HNF1AKD relative to control β cells. *P*= Benjamini-Hochberg adjusted *P*-value; all *P*<0.05.

Gene Ontology (GO) pathway analysis of genes downregulated after *HNF1A* loss identified known regulators of hormone secretion (*CHGA, CHGB, UCN3, PCSK2, TSPAN7, CACNA1D, TMEM27*), extracellular matrix (ECM) organization and cell adhesion (ADAMTS2, *COL6A2, COL1A1, COL4A2, CDH12*), cell-cell signaling (*GIPR, GLP1R, FFAR1, PRKCA, INSR*), glucose homeostasis (*G6PC2, PP1R1A, DEPTOR, ATP2A3, SLC2A2*), and endocrine pancreas development (*PAX4, MAFB, TM4SF4, FOXA2, ISL1, PAX6, NEUROD1*) (Fig. 3E). Additionally, downregulated genes were significantly enriched for KEGG pathways related to maturity onset diabetes of the young (*HNF1A, HNF4A, PAX4, INS, NEUROD1, GCK)* and cAMP signaling (*ADCY1, PDE3B, PFFAR2, FXYD2, GPR119*) (Fig. 3G).

By contrast, GO term analysis revealed that genes upregulated upon *HNF1A* loss were related to nucleic acid metabolic processes (*ATF3, FBXO4, TRIM11, DCP1A*), type I interferon responses (*IRF7, ISG20, ISG15, MX2*), and transcription repressor activity (*GZF1, NR1D1, HES1, HBP1, SCML1*) (Fig. 3F). These results suggest *HNF1A* is necessary to maintain the expression of hundreds of crucial genes encoding hallmark regulators of human β cell function.

### RNA-seq identifies adult human α cell transcriptome changes after *HNF1A* loss

Our evidence of α cell dysregulation after HNF1AKD (Fig. 1H-J and 2F-K) and prior studies by others (Haliyur et al. 2019; Sato et al. 2020) indicate that HNF1α is required to maintain adult α cell function. However, little is known about HNF1α gene regulation in mature human α cells, aside from studies of a single patient-derived sample (Haliyur et al. 2019). Here we used RNA-seq to investigate the *HNF1A*-dependent human α cell transcriptome. To isolate HNF1AKD α cells, we used FACS to enrich for transduced GFP^+^ cells expressing the human α cell marker CD26 (Augstein et al. 2015; Arda et al. 2016) (Fig. 4A). Glucagon (*GCG*) enrichment in the CD26^+^ cell fraction was verified by RT-qPCR (Supplemental Fig. S4A-B). Consistent with prior reports (Blodgett et al. 2015; Arda et al. 2016), analysis of RNA-seq libraries generated from α cell fractions revealed that *HNF1A* mRNA was higher in control α cells versus control β cells (transcripts per million, TPM; Fig. 4B; Fig. 3B). Similar to suppression of *HNF1A* in β cells, we achieved greater than 70% suppression of *HNF1A* in α cells using lenti-shRNA (Fig. 4B).

**Figure 4.**
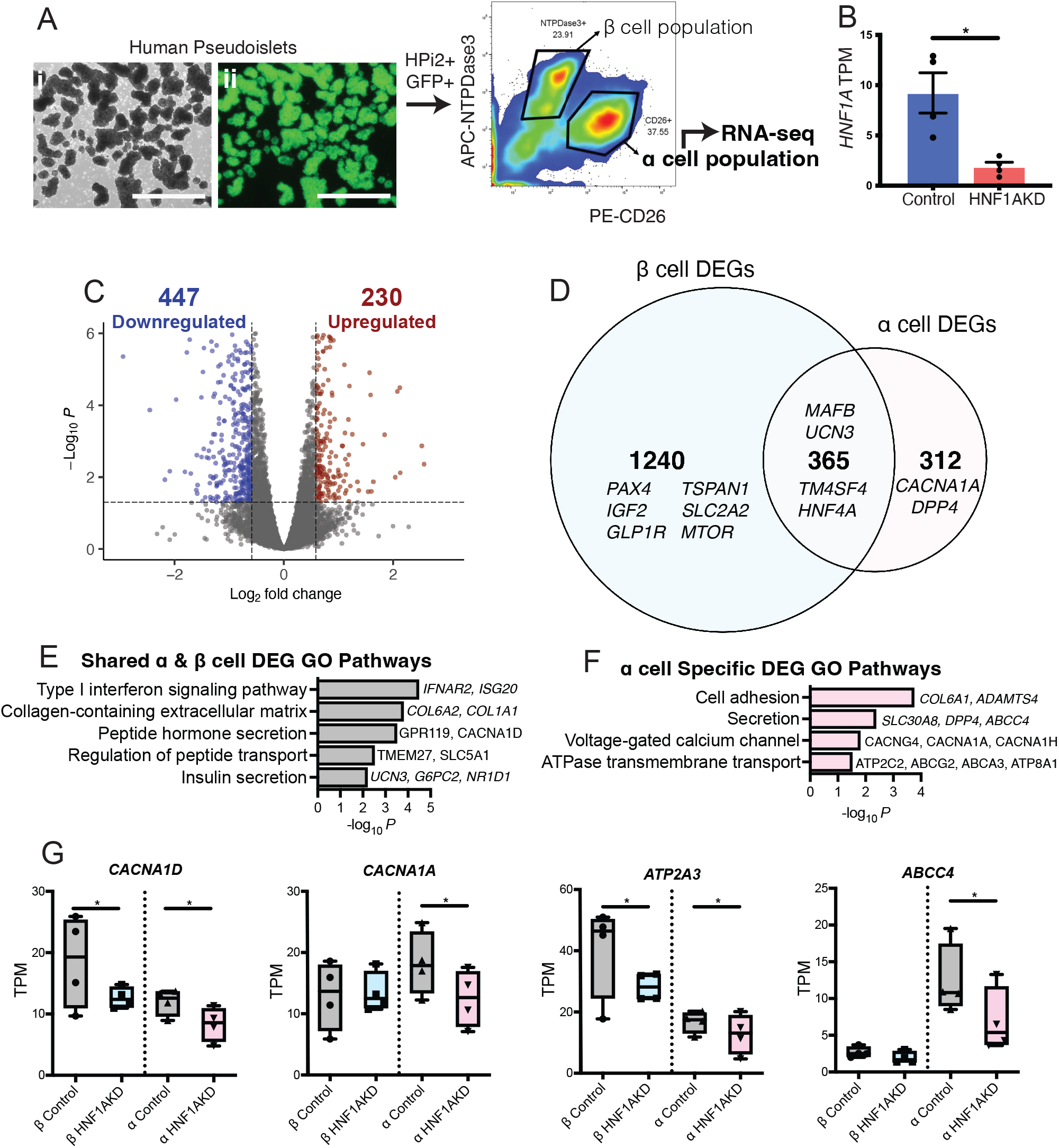
RNA-seq of HNF1AKD α cells identifies dysregulation of calcium channel complexes and ATPase-coupled transmembrane transport, as well as hormone secretion pathways shared with β cells. (A) Schematic of methods for isolation of transduced α cells (HPi2+GFP+CD26+) from control and HNF1AKD pseudoislets for downstream RNA sequencing; (i) brightfield and (ii) blue light (488nm) images of human HNF1AKD pseudoislets, scale bars=1000μm. (B) *HNF1A* transcripts per million (TPM) in sequenced α cell samples. (C) Differential expression (DE) analysis revealed significantly up- and down-regulated genes after HNF1AKD in α cells; thresholds: FC= 1.5, adjusted *P*-value= 0.05. (D) Venn diagram comparing α versus β cell DEGs revealed shared and cell-specific consequences of HNF1AKD. Gene ontology (GO) pathways of (E) shared and (F) α cell enriched DEG sets. (G) Boxplots displaying TPM of select DEGs. *P*= Benjamini-Hochberg adjusted *P*-value; \**P*<0.05; all *P*<0.05 for GO pathways.

DESeq2 analysis of HNF1AKD versus control RNA-seq libraries demonstrated 447 downregulated and 230 upregulated genes after *HNF1A* loss in α cells (Fig. 4C). More than half of α cell DEGs (365/677) were also dysregulated in β cells (Fig. 4D). These included genes encoding known developmental regulators and HNF1α targets (*TM4SF4, HNF4A*) and critical pancreatic islet regulators (*MAFB, UCN3*). GO term analysis of this ‘overlapping’ gene set highlighted shared pathways related to type I interferon signaling (*IFNAR2, ISG20*), collagen-containing ECM (*COL6A2, COL1A1*), and peptide hormone secretion (*CACNA1D, GPR119, TMEM27, SLC5A1, G6PC2*) (Fig. 4E).

GO term analysis of DEGs also identified α cell-specific changes after *HNF1A* loss, including enrichment of pathways related to cell adhesion (*COL6A1, ADAMTS4*) and hormone secretion, including the α cell-enriched factor *ABCC4* (Fig. 4F). Additional GO pathways enriched in α cell specific DEGs included voltage-gated Ca^2+^ channel constituents and ATPase-coupled transmembrane transport (Fig. 4F). While *CACNA1D* expression was significantly downregulated in both β and α cells after *HNF1A* loss, the expression of several additional voltage-gated Ca^2+^ channel genes (*CACNA1A, CACNG4, CACNA1H*) was changed in α cells but not in β cells (Fig. 4G). The *ATP2A3* gene encodes an enzyme that couples ATP hydrolysis with Ca^2+^ transport into the endoplasmic reticulum (ER), and was downregulated in both α and β cells after *HNF1A* suppression. However, additional ATP-coupled transporters (*ABCC4, ABCG2, ABCA3, ATP8A1*) were downregulated in α cells but not in β cells (Fig. 4G). Thus, our studies provide evidence for HNF1α direct or indirect regulation of hundreds of human pancreatic α cell genes.

### Direct targets of HNF1α identified by CUT&RUN

To identify direct HNF1α target genes in primary human islet cells, we performed CUT&RUN, a high-resolution assay capable of assessing transcription factor DNA binding sites *in situ* (Skene and Henikoff 2017). To overcome low endogenous islet expression of HNF1α and the reduced yields inherent to primary human pancreatic samples, we used lentiviral transduction to express a transgene encoding human *HNF1A* tagged with the FLAG immuno-epitope in human islet cells (Fig. 5A). Afterwards, DNA bound by HNF1α-FLAG protein was enriched with an anti-FLAG antibody and sequenced. This approach captured approximately 150bp fragments that encompassed HNF1α-FLAG-bound and immediately adjacent DNA regions. We used HOMER (Heinz et al. 2010) to identify genomic regions captured by HNF1α-FLAG (n=3 donors; cumulative Poisson *P*<0.0001 versus anti-IgG antibody control samples) as previously reported (Bevacqua et al. 2021b). We observed enriched read densities in HNF1α-FLAG CUT&RUN DNA peak centers compared to minimal enrichment at these sites for IgG control samples (Supplemental Fig. S5A-B). HOMER motif analysis identified that HNF1α-FLAG-bound genomic peaks were significantly enriched for the HNF1α binding motif (Fig. 5B), and other transcription factor motifs previously shown to be enriched in pancreatic islet enhancer clusters (Pasquali et al. 2014), including PDX1 (Fig. 5B) and NKX6.1 (Supplemental Fig. S5C).

**Figure 5.**
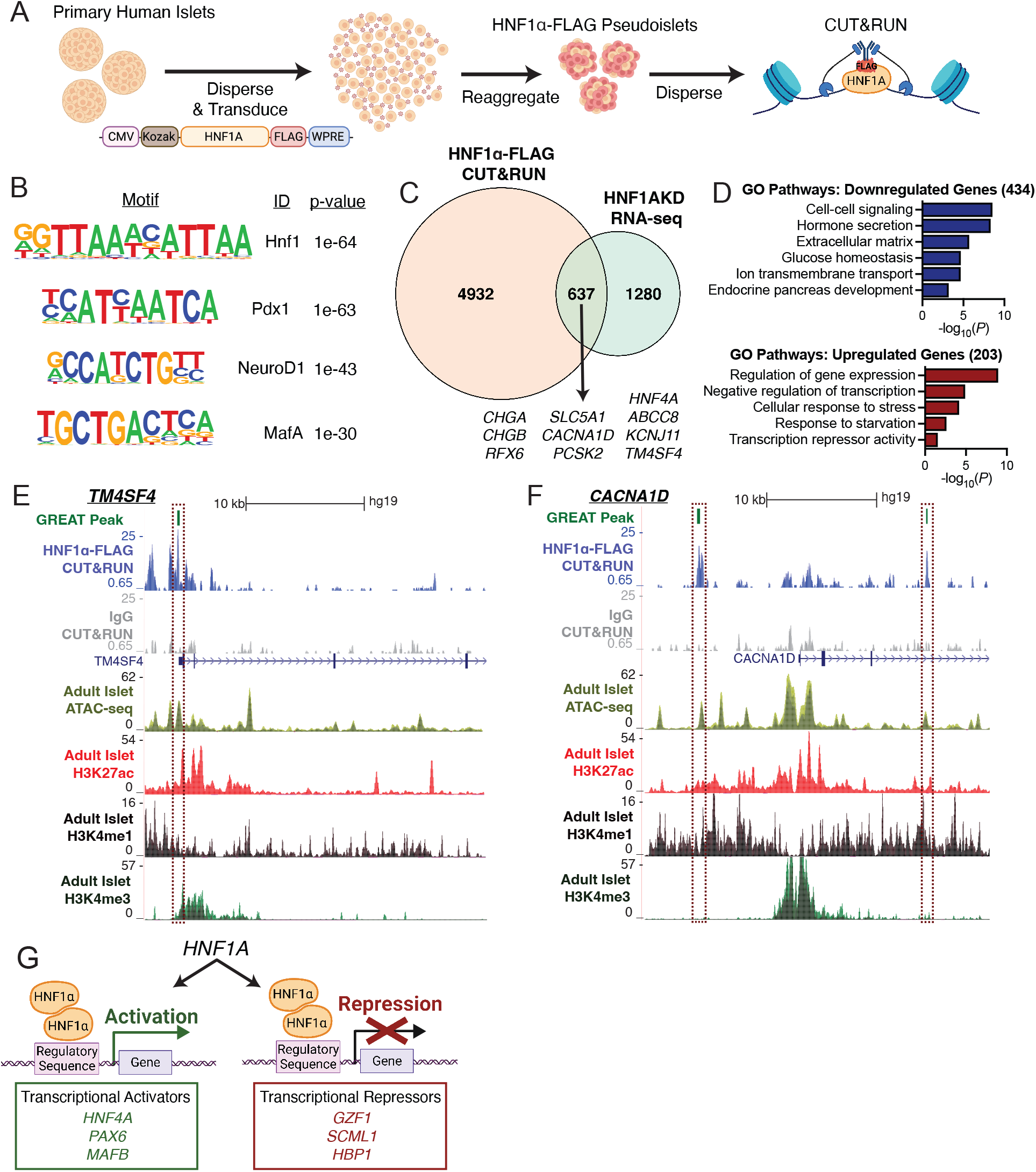
CUT&RUN identifies direct binding targets of HNF1α in primary human islet cells. (A) Schematic of methods: pseudoislets expressing HNF1α-FLAG were used for CUT&RUN with anti-FLAG antibody (n=3 donors). (B) Enriched motifs in the HNF1α-FLAG CUT&RUN peaks (versus IgG controls). (C) Venn diagram of genes associated with HNF1α-FLAG peaks identified by CUT&RUN (HNF1α-FLAG CUT&RUN) versus HNF1AKD differentially expressed genes in primary islet cells identified by RNA-seq (HNF1AKD RNA-seq). (D) Gene ontology (GO) pathway analysis of overlapping genes from panel C, subset into genes that were downregulated or upregulated in RNA-seq analysis. (E, F) UCSC Genome Browser tracks showing genomic regions associated with HNF1α-FLAG CUT&RUN peaks near the genes (E) *TM4SF4* and (F) *CACNA1D*; HNF1α-FLAG CUT&RUN enriched peaks identified by Genomic Regions Enrichment of Annotations Tool (GREAT) are highlighted in dashed boxes, and regulated genes are depicted below IgG control tracks. Accessible chromatin regions in human islets are shown by ATAC-seq and ChIP-seq (H3K427ac, H3K4me1, and H3Kme3) tracks (Pasquali et al. 2014). (G) Schematic depicting HNF1α’s dual role as a transcriptional activator and repressor in pancreatic islet cells.

Using the GREAT algorithm (McLean et al. 2010), we associated HNF1α-FLAG-bound regions to 5,569 proximate genes. To prioritize direct regulatory targets of HNF1α, we compared these genes to DEGs identified by RNA-seq after *HNF1A* loss. Of 1,917 DEGs in HNF1AKD α or β cells, 637 were also present in the HNF1α-FLAG CUT&RUN gene set (Fig. 5C). The concordance between HNF1α-FLAG-bound regions and DEGs after *HNF1A* loss provides evidence that our CUT&RUN approach identified direct HNF1α targets.

Consistent with reports that HNF1α functions as a transcriptional activator (Boj et al. 2010; Soutoglou et al. 2000; Luco et al. 2008), 68% (434/637) of the putative HNF1α targets identified by the intersection of our HNF1α-FLAG CUT&RUN and HNF1AKD RNA-seq gene sets were downregulated upon *HNF1A* loss (Fig. 5D). These downregulated genes were enriched for GO pathways related to cell-cell signaling *(CASR, DPP4, CADM1, MMP16, RGS4)*, hormone secretion (*ABCC8, KCNJ11, CHGA, CHGB, PCSK2*), ECM *(COL6A3, ADAMTS2, TGM2, TGFBI)*, glucose homeostasis *(G6PC2, GCK, SRI, FFAR2)*, ion transmembrane transport (*ATP2A3, CACNA1D, SLC5A1, SLC30A8, TMEM37*), and endocrine pancreas development (*HNF4A, TM4SF4, NEUROD1, PAX6, RFX6, MAFB*). As expected, HNF1α-FLAG-bound genomic regions localized to presumptive accessible promoter and enhancer regions, as revealed by colocalization with transposase integration sites and histone marks reported in prior ATAC-seq and ChIP-seq studies (Fig. 5E-F) (Mularoni et al. 2017; Pasquali et al. 2014). These findings support that HNF1α directly activates expression of target genes that encode factors essential for mature α and β cell function (Fig. 5G). By contrast, the 203 (32%) putative HNF1α targets upregulated after *HNF1A* loss were broadly related to negative gene regulation (*NR1D1, GZF1, HBP1, SCML1, BACH2, BHLHE41, E2F7*) (Fig. 5D). These findings implicate HNF1α as a direct negative regulator of transcriptional repressors, suggesting that HNF1α de-repression of transcriptional networks is another mechanism for maintaining human islet cell function (Fig. 5G).

### Islet transcriptomes after acute *HNF1A* loss and in congenital HNF1A-MODY

In subjects with HNF1A-MODY, diabetes typically occurs in adolescence or early adulthood. Thus, *HNF1A* haploinsufficiency manifests in these subjects after many years, and the mechanisms underlying a switch from sufficient islet cell function in childhood to subsequent islet insufficiency remain poorly understood. To assess the applicability of studying acute *HNF1A* suppression in human pseudoislets to understanding HNF1A-MODY, we compared HNF1α gene targets identified in this study with islet RNA-seq datasets from a subject with HNF1A-MODY (Haliyur et al. 2019).

The majority of HNF1α targets (368/637, 58%) we identified with CUT&RUN and RNA-Seq were differentially expressed in the RNA-seq data obtained from human HNF1A-MODY α and β cells. Pearson correlation analysis of normalized gene expression levels of these putative targets in our adult HNF1AKD α cells versus the HNF1A-MODY donor α cells revealed a highly significant positive correlation (r=0.42, *P*=2.2E-16). Similarly, we observed a positive correlation in gene expression levels of putative HNF1α targets between HNF1AKD β cells and the HNF1A-MODY donor β cells (r=0.32, *P=*5.3E-16). This concordance was further validated by heatmap visualization of dysregulated adult HNF1α targets in the HNF1A-MODY donor α cells (Fig. 6A) and β cells (Fig. 6B). Notably, many of the genes common to our dataset and the prior study were associated with ECM organization (*DOCK1, COL5A1, ADAM22*), ion transmembrane transport *(ATP2A3, FXYD2, SLC30A8)*, glucose metabolism (*SLC2A2, G6PC2, FFAR2*), hormone secretion (*ABCC8, KCNJ11, CACNA1D*), cell-cell communication (*CASR, DSCAML1, DPP4*), and transcriptional repression *(GZF1, SCML1, HBP1, PARD6G, ZBTB21, BACH2)*. Thus, our observations demonstrate an unexpectedly high concordance of DEGs in α and β cells of islets from a congenital HNF1A-MODY donor and from acute conditional *HNF1A* loss-of-function in primary human islets.

**Figure 6.**
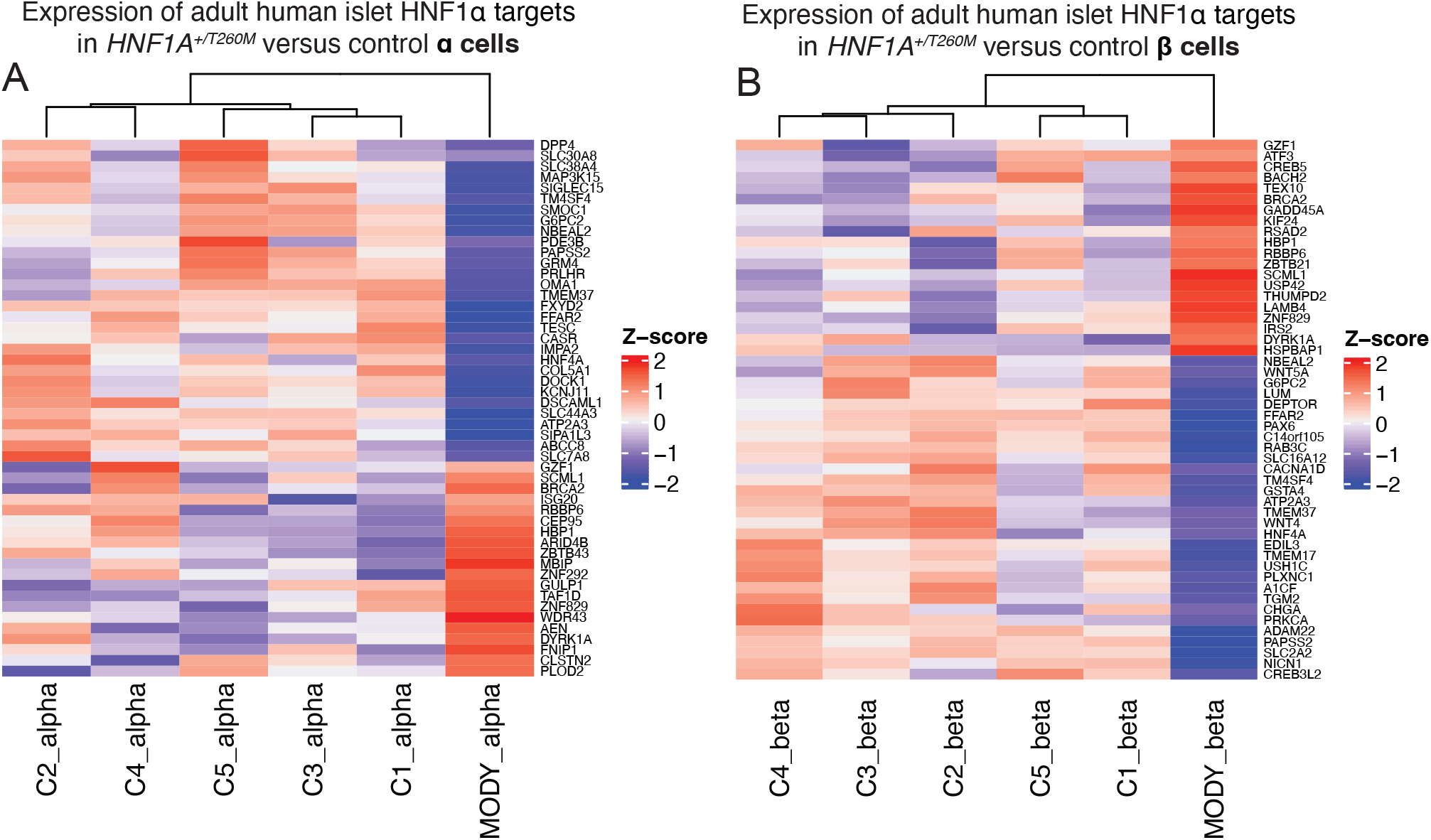
Comparison of HNF1α targets in primary human islets with HNF1A-MODY adult donor datasets demonstrates conserved HNF1α regulatory pathways that are critical for mature islet cell function. (A-B) Heatmaps showing relative expression of genes in (A) α and (B) β cells isolated from an *HNF1A*^*+/T260M*^ donor (MODY) versus healthy control donors (C1-C5); genes depicted were top differentially expressed genes in primary islet HNF1AKD RNA-seq data also identified in HNF1α-FLAG CUT&RUN data (putative adult HNF1α targets).

## DISCUSSION

Mutations in *HNF1A* account for the commonest form of human monogenic diabetes (Riddle et al. 2020), and *HNF1A* variants are also associated with T2D risk (Najmi et al. 2017; Althari et al. 2020). Here, we used pseudoislet-based genetics to achieve *HNF1A* loss-of-function in primary human islets. Through RNA sequencing, chromatin-based assays, and functional studies, we elucidated genetic targets and regulatory functions governed by HNF1α in mature human pancreatic α and β cells. This innovative strategy reconstituted islet phenotypes reported in patients with HNF1A-MODY, and our findings highlight the value of conditional genetic studies in adult human pseudoislets.

Previous reports have used stem-cell derived models of *HNF1A* deficiency to study the role of HNF1α during development of progeny resembling β cells (Cardenas-Diaz et al. 2019; Low et al. 2021; Cujba et al. 2022). Here, we acutely suppressed *HNF1A* in adult human islet cells, and observed transcriptional and functional phenotypes within days of *HNF1A* loss. Despite differences in the cell types studied and the timing of *HNF1A* loss, we found common features of gene regulation between prior stem-cell based studies and our data: both approaches revealed that HNF1α regulates genes critical for pancreas development, glucose metabolism and hormone secretion (like *HNF4A, TM4SF4, PAX4, PPP1R1A, G6PC2, SLC2A2, TMEM27, TMEM37, SLC38A4, GIPR, A1CF, ABCC8*, and *KCNJ11*). We also observed distinct features of gene regulation in our study. For example, *HNF1A* loss in human embryonic stem cell (hESC)-derived β-like cells decreased *PAX4* expression and increased expression of α cell ‘signature’ factors like *GCG* (Cardenas-Diaz et al. 2019; González et al. 2022). While we also observed decreased *PAX4* transcripts in HNF1AKD β cells, this was not accompanied by an increase in α cell-enriched genes. HNF1AKD in primary human islet cells also reduced expression of *MAFB, NEUROD1, PAX6, ISL1*, and *FOXA2*, transcription factors that are associated with both mature β and α cell functions (Lyttle et al. 2008). These changes were accompanied by blunted glucose-stimulated insulin secretion from HNF1AKD β cells and dysregulated glucagon secretion in HNF1AKD α cells. Thus, our findings indicate that HNF1α orchestrates genetic pathways crucial for hormone secretion in mature adult β and α cells.

There are clear glucagon secretion phenotypes in HNF1A-MODY patients (Haliyur et al. 2019; Tuomi et al. 2006; Østoft et al. 2014) and in rodents deficient for *Hnf1a* (Sato et al. 2020), but the basis for these findings was unknown, motivating our studies to elucidate HNF1α functions in human pancreatic α cells. Here, we characterized impaired α cell function after *HNF1A* loss, then linked physiological phenotypes with α cell transcriptional changes. Analysis of α cell gene expression after *HNF1A* loss highlighted multiple dysregulated pathways, including impaired regulation of Ca^2+^ channel subunit and ATPase-coupled transmembrane transport genes. Ca^2+^ influx is a well-established component of regulated islet cell hormone secretion (Klec et al. 2019), but previous studies in human models of *HNF1A* insufficiency have left the role of HNF1α in governing islet cell Ca^2+^ transients unresolved. In a recent study of stem cell-derived endocrine progenitors heterozygous for the HNF1A^H126D^ mutation, no defects in Ca^2+^ channel gene expression or Ca^2+^ responses to ionomycin challenge were observed (Low et al. 2021). However, another study in hESC-derived β-like cells reported that homozygous *HNF1A* knockout impaired a network of genes required for Ca^2+^ signaling (González et al. 2022). Similarly, a prior study reported significantly decreased *CACNA1H* mRNA levels in β cells and *CACNA1I, CACNA1A*, and *CACNA1H* mRNA levels in α cells from an adult HNF1A-MODY donor (Haliyur et al. 2019). Our work using primary human islets supports the view that HNF1α regulates voltage-gated Ca^2+^ channel subunit expression and Ca^2+^ channel currents in mature α cells.

Prior work based on β-like cells derived from stem cells has focused on *HNF1A* loss-of-function phenotypes in glucose uptake and ATP production (Low et al. 2021), as well as glycolysis and mitochondrial respiration (Cardenas-Diaz et al. 2019). Here, we report that other ATP-mediated pathways are also dysregulated upon *HNF1A* loss, both in β and α cells. *ATP2A3*, which encodes an enzyme that couples ATP hydrolysis with Ca^2+^ transport into the ER (Primeau et al. 2018), was downregulated after HNF1AKD in both α and β cells. In α cells, HNF1AKD resulted in reduced expression of several additional ATP transporters, including *ATP2C2*, which encodes an ATP-driven Golgi transporter of Ca^2+^, a key cofactor for processing and trafficking secretory pathway proteins (Vanoevelen et al. 2005). These findings suggest that *HNF1A* loss leads to global perturbation of ATP-associated processes in adult islet cells, and loss of ATP-driven Ca^2+^ transporters may aggravate anomalous glucagon secretion from α cells in *HNF1A*-deficient diabetes. This work also highlights the value of studying genetic mechanisms in native human islet cells.

Our work revealed significant overlap between genes dysregulated in primary α and β cells after *HNF1A* loss, including human pancreatic genes like *UCN3* and *MAFB* that regulate hormone secretion. Furthermore, our study provides evidence that HNF1α promotes normal ECM dynamics and suppresses interferon signaling in both cell types. This is consistent with work showing genes in these pathways are dysregulated in HNF1A-MODY; specifically, the interferon-associated genes *NLRC5, RSAD2*, and *SRPK1* were expressed at higher levels and the ECM-associated genes *COL5A1, COL1A2, COL1A1*, and *ADAM22* were expressed at lower levels in HNF1A-MODY α and β cells compared to healthy controls (Haliyur et al. 2019). ECM plays a critical role in pancreatic islet cell function and survival, and ECM addition to transplanted islets can enhance islet insulin secretion (Llacua et al. 2018). Additionally, accumulating evidence implicates activation of type I interferon signaling in the pathogenesis of T1D (Marroqui et al. 2021), and previous studies have demonstrated that HNF1α functions to suppress type I interferon signaling in the liver (He et al. 2022). The significant *concordance* of gene expression we observed with that reported in a case of HNF1A-MODY (Haliyur et al. 2019) further indicates that our model of acute *HNF1A* loss in primary human islet cells accurately recapitulated transcriptional phenotypes observed in human *HNF1A* insufficiency.

By correlating HNF1α-bound genomic regions determined through CUT&RUN with DEGs after *HNF1A* loss from our RNA-seq data, we also identified hundreds of genes likely to be directly regulated by HNF1α in adult human islets. These data verified HNF1α binding of putative target genes (including *HNF4A, TM4SF4, SLC2A2, KCNJ11, ABCC8*, and *SLC5A1*) reported in previous human and mouse studies that did not achieve α and β cell purification (Low et al. 2021; Boj et al. 2010; Odom et al. 2004; Servitja et al. 2009; Boj et al. 2001). Furthermore, our motif analyses provide evidence that HNF1α collaborates with other critical pancreatic transcription factors (e.g. *PDX1*) at enhancer cluster sites to regulate islet cell gene expression, consistent with prior findings (Pasquali et al. 2014).

While HNF1α is best known as a transcriptional activator that promotes expression of target genes (Low et al. 2021; Boj et al. 2010; Soutoglou et al. 2000; Luco et al. 2008), differential gene expression studies consistently demonstrate that HNF1α loss-of-function leads to both significant down-and up-regulation of genes (Cardenas-Diaz et al. 2019; Low et al. 2021; Haliyur et al. 2019; Servitja et al. 2009). Furthermore, direct transcriptional repression by HNF1α has been demonstrated in human and mouse hepatocytes (Patitucci et al. 2017). Thus, upregulation of genes following *HNF1A* loss could reflect indirect mechanisms of HNF1α-dependent transcriptional control (Piaggio et al. 1994; Kritis et al. 1993), like de-repression. Here, we report that approximately one-third of HNF1α target genes were upregulated after *HNF1A* loss; the majority of these were also upregulated in islets isolated from an HNF1A-MODY donor (Haliyur et al. 2019). Notably, many of these upregulated targets encode transcriptional repressors with previously characterized roles in modulating branching morphogenesis (e.g. GZF1) (Fukuda et al. 2003), or in restricting proliferation (HBP1) (Yee et al. 2004), homeotic gene expression (SCML1) (van de Vosse et al. 1998) and antioxidant response pathways (BACH2) (Son et al. 2021). These findings support our view that HNF1α has dual roles in islet cell transcriptional activation and de-repression of gene networks (Fig. 5G). Dynamic transcriptional de-repression is critical for multiple physiological processes: in addition to those noted above, these include circadian regulation (Chiou et al. 2016), inflammation (Perissi et al. 2010; Huang et al. 2009), and endocrine cell differentiation (Martin et al. 2015). Thus, diabetic islet phenotypes associated with *HNF1A* loss could reflect both loss of direct HNF1α-dependent transcriptional activation and aberrant transcriptional repression.

In summary, this study elucidates genomic targets of HNF1α in primary human islets and correlates loss of *HNF1A* with dysregulated gene expression and functional deficits in pancreatic α and β cells that phenocopy those in human *HNF1A*-deficient diabetes. We demonstrate that HNF1α directly activates genetic pathways crucial for regulated hormone secretion, and provide evidence that loss of HNF1α-dependent transcriptional repression could disrupt genetic pathways in islets with *HNF1A* deficiency. These findings advance our understanding of mechanisms underlying α and β cell defects in human monogenic diabetes and T2D.

## MATERIALS AND METHODS

### Human Islet Procurement

Deidentified human islets were obtained from nondiabetic organ donors procured through the Integrated Islet Distribution Network (IIDP), National Diabetes Research Institute (NDRI), International Institute for the Advancement of Medicine (IIAM), University of California San Francisco (UCSF), and the Alberta Diabetes Institute IsletCore (www.isletcore.ca). Supplemental Table S1 contains donor details.

### Constructs and Lentivirus Production

Lentiviral constructs coding for shRNA targeting exon 4 of human *HNF1A* were obtained from Dharmacon. plenti-CMV-HNF1A-cMyc-DDK was used in CUT&RUN experiments (OriGene Technologies RC211201L1). Lentiviruses were produced by transient transfection of HEK293T cells with lentiviral constructs, and with pMD2.G and psPAX2 packaging constructs (Addgene 12259 & 12260). TurboFect reagents were used for transfection (Thermo Scientific R0531). Supernatants were purified using PEG-it (System Biosciences); concentrated lentivirus was stored at −80°C.

### Human Pseudoislet Generation

Human islets were dispersed into single cells by enzymatic digestion (Accumax, Invitrogen) and transduced with 1×10^9^ viral units/mL lentivirus. Transduced cells were cultured in ultra-low attachment cell culture plates (Corning) for 5 days prior to analysis.

### Quantitative RT-PCR

RNA was isolated from whole pseudoislets using the PicoPure RNA isolation kit (Life Technologies). cDNA was synthesized using the Maxima first strand kit (Thermo Scientific), and gene expression was assessed by PCR using TaqMan gene expression mix (Thermo Scientific) and the following probes: ACTIN-B (Hs4352667_m1), HNF1A (Hs01551750_m1), HNF4A (Hs00604435_m1), HNF1A-AS1 (Hs00703760_s1), TMEM27 (Hs00252907_m1), KCNJ11 (Hs00265026_s1), SLC2A2 (Hs01096904_m1), SLC2A1 (Hs00892681_m1), SLC2A3 (Hs00359840_m1), INS (Hs00355773_m1), GCG (Hs01031536_m1).

### *In vitro* Insulin and Glucagon Secretion Assays

Batches of 30 pseudoislets were used for *in vitro* secretion assays as previously described (Peiris et al. 2018). Briefly, pseudoislets were incubated at 2.8mM then 16.7mM glucose or 6mM glucose then 1mM glucose+10mM L-arginine for 60min each, and supernatants were collected. Hormones in the supernatants and pseudoislet lysates were quantified using glucagon and human insulin ELISA kits (Mercodia).

### Patch-clamp electrophysiology studies

Single-cell patch-clamp studies were performed as described previously (Lin et al. 2021). Control and HNF1AKD human pseudoislets were dissociated to single cells and cultured in 5.5mM glucose media for 1-3 days. Prior to whole-cell patch-clamping, media was changed to a bath solution containing: 118mM NaCl, 20mM Tetraethylammonium-Cl, 5.6mM KCl, 1.2mM MgCl2, 2.6mM CaCl2, 5mM HEPES, and 5mM glucose (pH 7.4 with NaOH) in a heated chamber (32–35 °C). Patch-clamping was performed using fire polished thin wall borosilicate pipettes coated with Sylgard (3-5mOhm) and filled with the following intracellular solution: 125mM Cs-glutamate, 10mM CsCl, 10mM NaCl, 1mM MgCl2, 0.05mM EGTA, 5mM HEPES, 0.1mM cAMP, and 3mM MgATP (pH 7.15 with CsOH). Electrophysiological data were recorded using a HEKA EPC10 amplifier and PatchMaster Software (HEKA Instruments Inc, Lambrecht/Pfalz, Germany) within 5min of break-in. The stability of seal (>10 GOhm) and access resistance (<15 MOhm) throughout the experiment were assessed for quality control. FitMaster (HEKA Instruments Inc) was used for data analysis.

### Transplantation and *in vivo* assessment of pseudoislet function

Batches of 1000 pseudoislets were transferred into the renal capsular space of host animals using a glass micro-capillary tube. Transplant recipients were 3-month-old male NOD scid IL2Rγnull (“NSG,” The Jackson Laboratory 005557) or GKO-NSG (Tellez et al. 2020) mice. One month post transplantation, all mice received an intraperitoneal (IP) injection of 3g glucose/kg body weight. For sulfonylurea sensitivity testing, NSG mice received an injection of 2.5mg glibenclamide (Sigma-Aldrich G0639)/kg of body weight at 6 weeks post-transplant. For insulin tolerance tests, GKO mice received 0.5U Humulin R (Lilly)/kg body weight (in 0.9% NaCl) at 6 weeks post-transplantation. For all experiments, glucose measurements and blood samples for hormone quantification were collected via the tail vein. Blood glucose was measured using a glucometer (True Metrix), and hormones were measured using glucagon and human insulin ELISA kits (Mercodia).

### Extracellular Staining and FACS of human islet cells

Pseudoislets were dispersed into single cells, stained with the LIVE/DEAD Fixable Near-IR dead cell stain kit (Life Technologies), and washed with cell staining buffer (BioLegend 420201). Then, cells were stained with the following primary-secondary conjugated antibodies: HPi2-PE/Cy7 (Novus Biologicals NBP1-18946PECY7), NTPDase3-647 (clone hN3-B3S; Jean Sévigny Lab) and CD26-PE (BioLegend 302706). Labeled cells were sorted on a special order five-laser FACS Aria II (BD Biosciences) using a 100-μm nozzle, with appropriate compensation controls and doublet removal. Sorted cells were collected into low retention tubes containing 100μL of FACS buffer with RiboLock RNase inhibitor (Thermo Scientific).

### RNA-seq library preparation and data analysis

A total of ≥5,000 sorted live β or α cells were used for each RNA-seq library construction. RNA was isolated using the PicoPure RNA isolation kit (Life Technologies); input RNA for libraries had RIN >8. The SMART-seq v4 Ultra Low input RNA kit (Clontech) was used to amplify cDNA, and libraries were generated using the Nextera XT DNA Library Preparation Kit (Illumina). Barcoded libraries were sequenced as paired-end 150-bp (PE150) reads on the Illumina NovaSeq 6000 platform. All libraries had >30 million reads, and FastQC v0.11.9 was used for quality control. Barcodes were trimmed using Trim Galore v0.5.0. Reads were aligned to the human genome index (GRCh38, Ensembl release 104) using STAR v2.6.1d (Dobin et al. 2013). Estimated Counts and Transcripts per million (TPM) were quantified using RSEM v1.3.1 (Li and Dewey 2011). Differentially expressed genes with fold change were detected using the DESeq2 R package (Love et al. 2014) for HNF1AKD versus control conditions, controlling for donor differences. A *P*-adjusted cutoff of 0.05 and fold change threshold of 1.5 were used to filter significant DEGs. g:Profiler version e106_eg53_p16_65fcd97 was used for gene set enrichment analysis (Raudvere et al. 2019). RNA-seq datasets from HNF1A-MODY donor islets were obtained from Haliyur et al. (GEO: GSE106148, GSE116559, and GSE120299) (Haliyur et al. 2019). For comparison with the Haliyur et al. dataset, Pearson’s correlation analysis was performed on Z-transformed average HNF1AKD TPMs for each cell type (versus 4 control TPMs for β and α cell datasets) and Z-transformed values of the HNF1A-MODY donor (n=1 HNF1A-MODY donor versus 5 controls).

### CUT&RUN Assay and Library Preparation

CUT&RUN was performed on nuclei extracted from 500,000 dispersed HNF1α-FLAG pseudoislet cells per condition using CUTANA™ ChIC/ CUT&RUN protocol version 3.1. Briefly, nuclei were extracted with nuclear extraction buffer (20 mM HEPES–KOH at pH 7.9; 10 mM KCl; 0.1% Triton X-100; 20% glycerol; 1mM MnCl_2_; 0.5 mM spermidine; 1x Halt protease inhibitor cocktail, Thermo Scientific 78425) for 10 minutes on ice and immobilized onto Concanavalin-A beads (EpiCypher 21-1401). After blocking and washes, samples were incubated with 0.5 μg of rabbit anti-FLAG (Sigma-Millipore F7425) or rabbit anti-IgG (EpiCypher 13-0042) antibodies overnight at 4°C. pAG-MNase (EpiCypher 15-1016) was added to nuclei at a concentration of 1:20 and incubated at RT for 10min. Targeted chromatin digestion was induced by adding 100mM CaCl_2_ and nutating samples for 2hrs at 4°C. DNA fragments were released and purified using the CUTANA™ ChIC/ CUT&RUN kit, according to the manufacturer’s instructions. DNA was resuspended in 0.1X TE and used for library preparation with the CUTANA™CUT&RUN Library Prep Kit (Epicypher 14-1001), according to the v1 manual. Libraries were sequenced as PE150 reads on the NovaSeq platform.

### CUT&RUN Data Analysis

All CUT&RUN libraries had >25 million reads. Reads were trimmed and aligned using CUT&RUNtools (Zhu et al. 2019). Trimmomatic was used for trimming (Bolger et al. 2014), Bowtie2 for alignment (Langmead and Salzberg 2012), and HOMER for peak calling using macs2.narrow outputs (Heinz et al. 2010). *P*-values for motif enrichment were generated by HOMER software. Genome browser tracks were generated from mapped reads using the “makeUCSCfile” command. The GREAT algorithm was used for gene annotation, using default parameters (McLean et al. 2010).

### Other Statistical Analysis

For RT-qPCR, GSIS, AUC and immunostaining quantification, the number of biological replicates, measure of central tendency and deviation, and statistical test used for analysis are detailed in each figure legend. Graphs and statistical analysis were produced using GraphPad Prism (v9) and R (v4.1.1) software.

## Data Visualization

Cytometry data were analyzed and graphed using FlowJo software (v10.8). Venn Diagrams and heat maps were generated in R using VennDiagram and ComplexHeatmap. Browser tracks were generated using the UCSC genome browser (Kent et al. 2002) and method graphics were created with BioRender.com.

## Data Availability

The raw sequencing data from this publication will be deposited in NCBI’s GeneExpression Omnibus (Edgar et al. 2002).

## ACKNOWLEDGMENTS

We thank Drs. R. Whitener, Y. Hang, J. Qian, K. Tellez, H. Peiris, M. Miranda, Mr. J. Lam and Mr. R. Rodriguez for technical assistance and advice. We also thank Professors A. Gloyn, R. Nusse, D. Kingsley, and Dr. N. Krentz for advice and encouragement. We gratefully acknowledge organ donors and their families, and the Alberta Diabetes Institute IsletCore, Integrated Islet Distribution Program (NIH UC4 DK098085), National Disease Research Interchange, and International Institute for the Advancement of Medicine. In Alberta, we thank the Human Organ Procurement and Exchange (HOPE) program and Trillium Gift of Life Network (TGLN), and James Lyon for his efforts in human islet isolation. RNA-seq data were generated using instrumentation purchased with NIH funds (S10OD025212 and 1S10OD021763). M.Q. is a student in the Stanford MSTP and Knight-Hennessey Scholars Program, and was previously supported by the Stanford Medical Scholars and HHMI Medical Research Fellows Programs. V.N. was supported by a NIH T32 training grant (5T32GM007790). R.B. was supported by a postdoctoral fellowship from JDRF (3-PDF-2018-584-A-N). Work here was supported by NIH awards (R01 DK107507; R01 DK108817; U01 DK123743; R01 DK126482 to S.K.K. and P.E.M.), and JDRF Center of Excellence (to S.K.K. and M. Hebrok). Work here was also supported by NIH P30 DK116074 (S.K.K.), which supports the Stanford Islet Research Core and Diabetes Genomics and Analysis Core.

## AUTHOR CONTRIBUTIONS

M.F.Q. and S.K.K. conceptualized the study. M.F.Q., R.J.B., V.M.N., X.L., W.Z., C.C., X.G. and X.D. performed experiments. M.F.Q. and S.K.K. wrote the manuscript with input from all coauthors. S.K.K. supervised the study. M.F.Q., R.J.B, V.M.N., P.E.M., and S.K.K. acquired the funding.

## REFERENCES

Althari S, Najmi LA, Bennett AJ, Aukrust I, Rundle JK, Colclough K, Molnes J, Kaci A, Nawaz S, van der Lugt T, et al. 2020. Unsupervised Clustering of Missense Variants in HNF1A Using Multidimensional Functional Data Aids Clinical Interpretation. Am J Hum Genet 107: 670–682.

Arda HE, Li L, Tsai J, Torre EA, Rosli Y, Peiris H, Spitale RC, Dai C, Gu X, Qu K, et al. 2016. Age-Dependent Pancreatic Gene Regulation Reveals Mechanisms Governing Human β Cell Function. Cell Metab 23: 909–920.

Augstein P, Naselli G, Loudovaris T, Hawthorne WJ, Campbell P, Bandala-Sanchez E, Rogers K, Heinke P, Thomas HE, Kay TW, et al. 2015. Localization of dipeptidyl peptidase-4 (CD26) to human pancreatic ducts and islet alpha cells. Diabetes Res Clin Pract 110: 291–300.

Bacon S, Kyithar MP, Rizvi SR, Donnelly E, McCarthy A, Burke M, Colclough K, Ellard S, Byrne MM. 2016. Successful maintenance on sulphonylurea therapy and low diabetes complication rates in a HNF1A-MODY cohort. Diabet Med J Br Diabet Assoc 33: 976–984.

Bevacqua RJ, Dai X, Lam JY, Gu X, Friedlander MSH, Tellez K, Miguel-Escalada I, Bonàs-Guarch S, Atla G, Zhao W, et al. 2021a. CRISPR-based genome editing in primary human pancreatic islet cells. Nat Commun 12: 2397.

Bevacqua RJ, Lam JY, Peiris H, Whitener RL, Kim S, Gu X, Friedlander MSH, Kim SK. 2021b. SIX2 and SIX3 coordinately regulate functional maturity and fate of human pancreatic β cells. Genes Dev 35: 234–249.

Blodgett DM, Nowosielska A, Afik S, Pechhold S, Cura AJ, Kennedy NJ, Kim S, Kucukural A, Davis RJ, Kent SC, et al. 2015. Novel Observations From Next-Generation RNA Sequencing of Highly Purified Human Adult and Fetal Islet Cell Subsets. Diabetes 64: 3172–3181.

Boj SF, Parrizas M, Maestro MA, Ferrer J. 2001. A transcription factor regulatory circuit in differentiated pancreatic cells. Proc Natl Acad Sci U S A 98: 14481–14486.

Boj SF, Petrov D, Ferrer J. 2010. Epistasis of Transcriptomes Reveals Synergism between Transcriptional Activators Hnf1α and Hnf4α. PLoS Genet 6: e1000970.

Bolger AM, Lohse M, Usadel B. 2014. Trimmomatic: a flexible trimmer for Illumina sequence data. Bioinforma Oxf Engl 30: 2114–2120.

Cardenas-Diaz FL, Osorio-Quintero C, Diaz-Miranda MA, Kishore S, Leavens K, Jobaliya C, Stanescu D, Ortiz-Gonzalez X, Yoon C, Chen CS, et al. 2019. Modeling Monogenic Diabetes using Human ESCs Reveals Developmental and Metabolic Deficiencies Caused by Mutations in HNF1A. Cell Stem Cell 25: 273-289.e5.

Chiou Y-Y, Yang Y, Rashid N, Ye R, Selby CP, Sancar A. 2016. Mammalian Period represses and derepresses transcription by displacing CLOCK-BMAL1 from promoters in a Cryptochrome-dependent manner. Proc Natl Acad Sci U S A 113: E6072–E6079.

Cujba A-M, Alvarez-Fallas ME, Pedraza-Arevalo S, Laddach A, Shepherd MH, Hattersley AT, Watt FM, Sancho R. 2022. An HNF1α truncation associated with maturity-onset diabetes of the young impairs pancreatic progenitor differentiation by antagonizing HNF1β function. Cell Rep 38: 110425.

Dai X-Q, Camunas-Soler J, Briant LJB, Dos Santos T, Spigelman AF, Walker EM, Arrojo E Drigo R, Bautista A, Jones RC, Avrahami D, et al. 2022. Heterogenous impairment of α cell function in type 2 diabetes is linked to cell maturation state. Cell Metab 34: 256-268.e5.

Dobin A, Davis CA, Schlesinger F, Drenkow J, Zaleski C, Jha S, Batut P, Chaisson M, Gingeras TR. 2013. STAR: ultrafast universal RNA-seq aligner. Bioinforma Oxf Engl 29: 15–21.

Edgar R, Domrachev M, Lash AE. 2002. Gene Expression Omnibus: NCBI gene expression and hybridization array data repository. Nucleic Acids Res 30: 207–210.

Enge M, Arda HE, Mignardi M, Beausang J, Bottino R, Kim SK, Quake SR. 2017. Single-Cell Analysis of Human Pancreas Reveals Transcriptional Signatures of Aging and Somatic Mutation Patterns. Cell 171: 321-330.e14.

Friedlander MSH, Nguyen VM, Kim SK, Bevacqua RJ. 2021. Pancreatic Pseudoislets: An Organoid Archetype for Metabolism Research. Diabetes 70: 1051–1060.

Fukuda N, Ichihara M, Morinaga T, Kawai K, Hayashi H, Murakumo Y, Matsuo S, Takahashi M. 2003. Identification of a novel glial cell line-derived neurotrophic factor-inducible gene required for renal branching morphogenesis. J Biol Chem 278: 50386–50392.

González BJ, Zhao H, Niu J, Williams DJ, Lee J, Goulbourne CN, Xing Y, Wang Y, Oberholzer J, Blumenkrantz MH, et al. 2022. Reduced calcium levels and accumulation of abnormal insulin granules in stem cell models of HNF1A deficiency. Commun Biol 5: 779.

Gromada J, Chabosseau P, Rutter GA. 2018. The α-cell in diabetes mellitus. Nat Rev Endocrinol 14: 694–704.

Haliyur R, Tong X, Sanyoura M, Shrestha S, Lindner J, Saunders DC, Aramandla R, Poffenberger G, Redick SD, Bottino R, et al. 2019. Human islets expressing HNF1A variant have defective β cell transcriptional regulatory networks. J Clin Invest 129: 246–251.

He J, Du C, Peng X, Hong W, Qiu D, Qiu X, Zhang X, Qin Y, Zhang Q. 2022. Hepatocyte nuclear factor 1A suppresses innate immune response by inducing degradation of TBK1 to inhibit steatohepatitis. Genes Dis.

Heinz S, Benner C, Spann N, Bertolino E, Lin YC, Laslo P, Cheng JX, Murre C, Singh H, Glass CK. 2010. Simple combinations of lineage-determining transcription factors prime cis-regulatory elements required for macrophage and B cell identities. Mol Cell 38: 576–589.

Huang W, Ghisletti S, Perissi V, Rosenfeld MG, Glass CK. 2009. Transcriptional integration of TLR2 and TLR4 signaling at the NCoR derepression checkpoint. Mol Cell 35: 48–57.

Kent WJ, Sugnet CW, Furey TS, Roskin KM, Pringle TH, Zahler AM, Haussler D. 2002. The human genome browser at UCSC. Genome Res 12: 996–1006.

Klec C, Ziomek G, Pichler M, Malli R, Graier WF. 2019. Calcium Signaling in ß-cell Physiology and Pathology: A Revisit. Int J Mol Sci 20: 6110.

Kritis AA, Ktistaki E, Barda D, Zannis VI, Talianidis L. 1993. An indirect negative autoregulatory mechanism involved in hepatocyte nuclear factor-1 gene expression. Nucleic Acids Res 21: 5882–5889.

Langmead B, Salzberg SL. 2012. Fast gapped-read alignment with Bowtie 2. Nat Methods 9: 357–359.

Li B, Dewey CN. 2011. RSEM: accurate transcript quantification from RNA-Seq data with or without a reference genome. BMC Bioinformatics 12: 323.

Lin H, Smith N, Spigelman AF, Suzuki K, Ferdaoussi M, Alghamdi TA, Lewandowski SL, Jin Y, Bautista A, Wang YW, et al. 2021. β-Cell Knockout of SENP1 Reduces Responses to Incretins and Worsens Oral Glucose Tolerance in High-Fat Diet-Fed Mice. Diabetes 70: 2626–2638.

Llacua LA, Faas MM, de Vos P. 2018. Extracellular matrix molecules and their potential contribution to the function of transplanted pancreatic islets. Diabetologia 61: 1261–1272.

Love MI, Huber W, Anders S. 2014. Moderated estimation of fold change and dispersion for RNA-seq data with DESeq2. Genome Biol 15: 550.

Low BSJ, Lim CS, Ding SSL, Tan YS, Ng NHJ, Krishnan VG, Ang SF, Neo CWY, Verma CS, Hoon S, et al. 2021. Decreased GLUT2 and glucose uptake contribute to insulin secretion defects in MODY3/HNF1A hiPSC-derived mutant β cells. Nat Commun 12: 3133.

Luco RF, Maestro MA, Sadoni N, Zink D, Ferrer J. 2008. Targeted deficiency of the transcriptional activator Hnf1alpha alters subnuclear positioning of its genomic targets. PLoS Genet 4: e1000079.

Lyttle BM, Li J, Krishnamurthy M, Fellows F, Wheeler MB, Goodyer CG, Wang R. 2008. Transcription factor expression in the developing human fetal endocrine pancreas. Diabetologia 51: 1169–1180.

Mahajan A, Taliun D, Thurner M, Robertson NR, Torres JM, Rayner NW, Payne AJ, Steinthorsdottir V, Scott RA, Grarup N, et al. 2018. Fine-mapping type 2 diabetes loci to single-variant resolution using high-density imputation and islet-specific epigenome maps. Nat Genet 50: 1505–1513.

Marroqui L, Perez-Serna AA, Babiloni-Chust I, Dos Santos RS. 2021. Chapter One - Type I interferons as key players in pancreatic β-cell dysfunction in type 1 diabetes. In International Review of Cell and Molecular Biology (eds. I. Santin and L. Galluzzi), Vol. 359 of Pancreatic ß-Cell Biology in Health and Disease, pp. 1–80, Academic Press.

Martin D, Kim Y-H, Sever D, Mao C-A, Haefliger J-A, Grapin-Botton A. 2015. REST represses a subset of the pancreatic endocrine differentiation program. Dev Biol 405: 316–327.

McLean CY, Bristor D, Hiller M, Clarke SL, Schaar BT, Lowe CB, Wenger AM, Bejerano G. 2010. GREAT improves functional interpretation of cis-regulatory regions. Nat Biotechnol 28: 495–501.

Miyachi Y, Miyazawa T, Ogawa Y. 2022. HNF1A Mutations and Beta Cell Dysfunction in Diabetes. Int J Mol Sci 23: 3222.

Mularoni L, Ramos-Rodríguez M, Pasquali L. 2017. The Pancreatic Islet Regulome Browser. Front Genet 8: 13.

Najmi LA, Aukrust I, Flannick J, Molnes J, Burtt N, Molven A, Groop L, Altshuler D, Johansson S, Bjørkhaug L, et al. 2017. Functional Investigations of HNF1A Identify Rare Variants as Risk Factors for Type 2 Diabetes in the General Population. Diabetes 66: 335–346.

Odom DT, Zizlsperger N, Gordon DB, Bell GW, Rinaldi NJ, Murray HL, Volkert TL, Schreiber J, Rolfe PA, Gifford DK, et al. 2004. Control of pancreas and liver gene expression by HNF transcription factors. Science 303: 1378–1381.

Østoft SH, Bagger JI, Hansen T, Hartmann B, Pedersen O, Holst JJ, Knop FK, Vilsbøll T. 2015. Postprandial incretin and islet hormone responses and dipeptidyl-peptidase 4 enzymatic activity in patients with maturity onset diabetes of the young. Eur J Endocrinol 173: 205–215.

Østoft SH, Bagger JI, Hansen T, Pedersen O, Holst JJ, Knop FK, Vilsbøll T. 2014. Incretin Effect and Glucagon Responses to Oral and Intravenous Glucose in Patients With Maturity-Onset Diabetes of the Young—Type 2 and Type 3. Diabetes 63: 2838–2844.

Pasquali L, Gaulton KJ, Rodríguez-Seguí SA, Mularoni L, Miguel-Escalada I, Akerman I, Tena JJ, Morán I, Gómez-Marín C, van de Bunt M, et al. 2014. Pancreatic islet enhancer clusters enriched in type 2 diabetes risk-associated variants. Nat Genet 46: 136–143.

Patitucci C, Couchy G, Bagattin A, Cañeque T, de Reyniès A, Scoazec J-Y, Rodriguez R, Pontoglio M, Zucman-Rossi J, Pende M, et al. 2017. Hepatocyte nuclear factor 1α suppresses steatosis-associated liver cancer by inhibiting PPARγ transcription. J Clin Invest 127: 1873–1888.

Peiris H, Park S, Louis S, Gu X, Lam JY, Asplund O, Ippolito GC, Bottino R, Groop L, Tucker H, et al. 2018. Discovering human diabetes-risk gene function with genetics and physiological assays. Nat Commun 9: 3855.

Perissi V, Jepsen K, Glass CK, Rosenfeld MG. 2010. Deconstructing repression: evolving models of corepressor action. Nat Rev Genet 11: 109–123.

Piaggio G, Tomei L, Toniatti C, De Francesco R, Gerstner J, Cortese R. 1994. LFB1/HNF1 acts as a repressor of its own transcription. Nucleic Acids Res 22: 4284–4290.

Pontoglio M, Sreenan S, Roe M, Pugh W, Ostrega D, Doyen A, Pick AJ, Baldwin A, Velho G, Froguel P, et al. 1998. Defective insulin secretion in hepatocyte nuclear factor 1alpha-deficient mice. J Clin Invest 101: 2215–2222.

Primeau JO, Armanious GP, Fisher ME, Young HS. 2018. The SarcoEndoplasmic Reticulum Calcium ATPase. Subcell Biochem 87: 229–258.

Raudvere U, Kolberg L, Kuzmin I, Arak T, Adler P, Peterson H, Vilo J. 2019. g:Profiler: a web server for functional enrichment analysis and conversions of gene lists (2019 update). Nucleic Acids Res 47: W191–W198.

Riddle MC, Philipson LH, Rich SS, Carlsson A, Franks PW, Greeley SAW, Nolan JJ, Pearson ER, Zeitler PS, Hattersley AT. 2020. Monogenic Diabetes: From Genetic Insights to Population-Based Precision in Care. Reflections From a Diabetes Care Editors’ Expert Forum. Diabetes Care 43: 3117–3128.

Sato Y, Rahman MM, Haneda M, Tsuyama T, Mizumoto T, Yoshizawa T, Kitamura T, Gonzalez FJ, Yamamura K-I, Yamagata K. 2020. HNF1α controls glucagon secretion in pancreatic α-cells through modulation of SGLT1. Biochim Biophys Acta Mol Basis Dis 1866: 165898.

Saunders DC, Brissova M, Phillips N, Shrestha S, Walker JT, Aramandla R, Poffenberger G, Flaherty DK, Weller KP, Pelletier J, et al. 2019. Ectonucleoside triphosphate diphosphohydrolase-3 antibody targets adult human pancreatic β-cells for in vitro and in vivo analysis. Cell Metab 29: 745-754.e4.

Servitja J-M, Pignatelli M, Maestro MA, Cardalda C, Boj SF, Lozano J, Blanco E, Lafuente A, McCarthy MI, Sumoy L, et al. 2009. Hnf1alpha (MODY3) controls tissue-specific transcriptional programs and exerts opposed effects on cell growth in pancreatic islets and liver. Mol Cell Biol 29: 2945–2959.

Skene PJ, Henikoff S. 2017. An efficient targeted nuclease strategy for high-resolution mapping of DNA binding sites. eLife 6: e21856.

Son J, Ding H, Farb TB, Efanov AM, Sun J, Gore JL, Syed SK, Lei Z, Wang Q, Accili D, et al. 2021. BACH2 inhibition reverses β cell failure in type 2 diabetes models. J Clin Invest 131: e153876.

Soutoglou E, Papafotiou G, Katrakili N, Talianidis I. 2000. Transcriptional activation by hepatocyte nuclear factor-1 requires synergism between multiple coactivator proteins. J Biol Chem 275: 12515–12520.

Tellez K, Hang Y, Gu X, Chang CA, Stein RW, Kim SK. 2020. In vivo studies of glucagon secretion by human islets transplanted in mice. Nat Metab 2: 547–557.

Tuomi T, Honkanen EH, Isomaa B, Sarelin L, Groop LC. 2006. Improved Prandial Glucose Control With Lower Risk of Hypoglycemia With Nateglinide Than With Glibenclamide in Patients With Maturity-Onset Diabetes of the Young Type 3. Diabetes Care 29: 189–194.

van de Vosse E, Walpole SM, Nicolaou A, van der Bent P, Cahn A, Vaudin M, Ross MT, Durham J, Pavitt R, Wilkinson J, et al. 1998. Characterization of SCML1, a new gene in Xp22, with homology to developmental polycomb genes. Genomics 49: 96–102.

Vanoevelen J, Dode L, Van Baelen K, Fairclough RJ, Missiaen L, Raeymaekers L, Wuytack F. 2005. The secretory pathway Ca2+/Mn2+-ATPase 2 is a Golgi-localized pump with high affinity for Ca2+ ions. J Biol Chem 280: 22800–22808.

Yee AS, Paulson EK, McDevitt MA, Rieger-Christ K, Summerhayes I, Berasi SP, Kim J, Huang C-Y, Zhang X. 2004. The HBP1 transcriptional repressor and the p38 MAP kinase: unlikely partners in G1 regulation and tumor suppression. Gene 336: 1–13.

Zhang H, Colclough K, Gloyn AL, Pollin TI. 2021. Monogenic diabetes: a gateway to precision medicine in diabetes. J Clin Invest 131: 142244.

Zhu Q, Liu N, Orkin SH, Yuan G-C. 2019. CUT&RUNTools: a flexible pipeline for CUT&RUN processing and footprint analysis. Genome Biol 20: 192.

