## Supplemental for "HNF1α transcriptional activation and repression maintain human islet α and β cell function"

### SUPPLEMENTAL MATERIALS AND METHODS

#### Immunofluorescence staining

Human pseudoislets (day 5) were fixed in 4% paraformaldehyde for 1hr at 4°C and embedded in collagen (Wako Chemicals). Mouse kidneys containing recovered human pseudoislet grafts were fixed in 4% paraformaldehyde for 1.5hrs at 4°C. All samples were dehydrated with 30% sucrose and embedded in OCT. Six-µm thick frozen sections were mounted onto slides and stained using standard cryostaining procedures. For immunostaining, sections were washed in PBS, permeabilized for 10 minutes using 0.2% Triton X, and incubated with blocking solution (5% normal donkey serum and 0.1% Triton X in PBS) for 1hr before overnight incubation with primary antibodies in blocking buffer at 4°C. The following primary antibodies were used: guinea pig anti-Insulin (1:200; Dako A0564), guinea pig anti-glucagon (1:2000; Takara M182), and rabbit anti-HNF1A (1:1000, Abcam ab204306). Slides were washed with PBS, incubated with Alexa Flour-conjugated secondary antibodies (488, 555, or 647; 1:500, donkey-anti-primary-host, Jackson ImmunoResearch) at room temperature for 1hr, incubated with Hoechst 33342 (Thermo Fisher Scientific, 1:2000) for 5min to detect nuclei, and preserved with mounting medium (Vector Labs, Vectashield H-1200). Images were obtained using a Zeiss AxioM1 microscope and a Leica SP2 confocal microscope. The ImageJ plug-in EzColocolization (Stauffer et al. 2018) was used for quantification of $GFP^{+}HNF1A^{+}$ cells from grafts.

#### References

Stauffer W, Sheng H, Lim HN. 2018. EzColocalization: An ImageJ plugin for visualizing and measuring colocalization in cells and organisms. *Sci Rep* **8**: 15764.

### SUPPLEMENTAL TABLES AND FIGURES

**Table S1:** Human Islet Donor Information

| Source | Sample ID | Age | Sex | BMI | HbA1c |
|---|---|---|---|---|---|
| IIDP | SAMN20478103 | 30 | Male | 25.4 | 5.8% |
| UCSF | rHIP148 | 38 | Female | 32.6 | 5.4% |
| Alberta IsletCore | R412 | 42 | Male | 29.9 | 4.8% |
| IIAM | SIRCPilot4 | 15 | Female | 22.9 | 5.3% |
| Alberta IsletCore | R440 | 35 | Female | 26.7 | 3.8% |
| UCSF | rHIP152 | 32 | Female | 29.2 | 5.0% |
| IIDP | SAMN28673685 | 37 | Male | 28.4 | 5.3% |
| IIAM | SIRC-CC | 18 | Female | 35 | Nondiabetic |
| IIAM | SIRC-1 | 44 | Female | 23.8 | Nondiabetic |
| IIDP | SAMN18092805 | 56 | Male | 21.6 | 5.1% |
| IIAM | SIRC-3 | 16 | Male | 35 | 5.5% |
| UCSF | rHIP150 | 36 | Female | 27.9 | 5.6% |
| IIDP | SAMN28673685 | 37 | Male | 28.4 | 5.3% |
| IIDP | SAMN28867622 | 36 | Male | 29.6 | 5.4% |
| Alberta IsletCore | R352 | 64 | Female | 20.4 | 5.3% |
| Alberta IsletCore | R357 | 64 | Male | 24.3 | 5.1% |
| IIDP | SAMN14132340 | 31 | Male | 27 | 5.2% |
| UCSF | rHIP141 | 41 | Male | 25.2 | 5.5% |
| IIDP | SAMN15770453 | 48 | Female | 30.9 | 5.8% |
| Alberta IsletCore | R421 | 60 | Female | 25.9 | Nondiabetic |
| UCSF | rHIP144 | 47 | Female | 23.5 | 5.7% |
| Alberta IsletCore | R373 | 31 | Male | 27.5 | 4.4% |
| Alberta IsletCore | R450 | 66 | Female | 21.9 | 4.3% |
| Alberta IsletCore | R447 | 62 | Male | 30.2 | 5.7% |

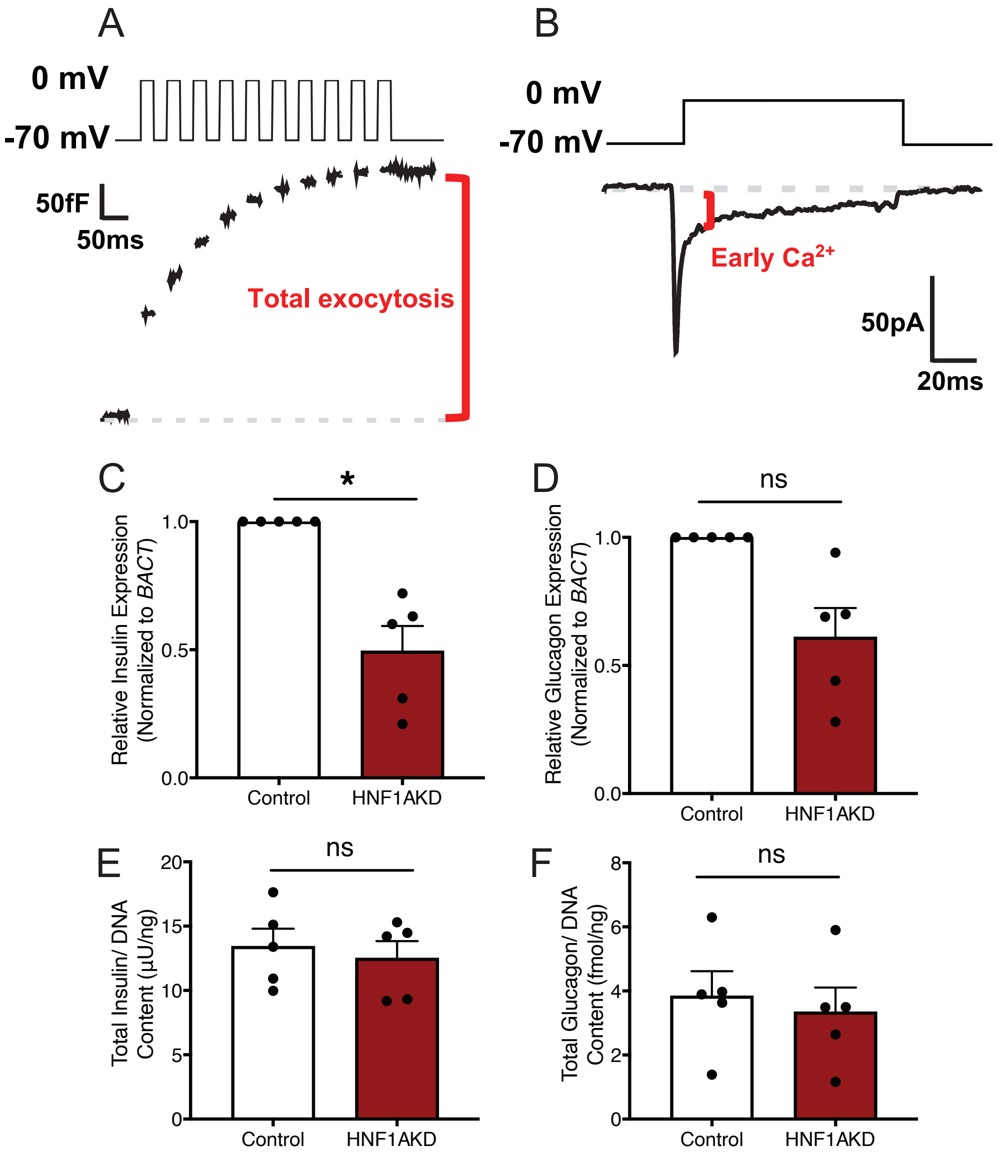


**Figure S1.** *In vitro* assessment of pseudoislet function and hormone content. Patch-clamp electrophysiology traces of (A) membrane capacitance and (B) current recordings from islet cells. (C) Insulin and (D) glucagon mRNA expression and total (E) insulin and (F) glucagon protein content in primary human pseudoislets 5 days after transduction with lenti-Control-shRNA ("Control") or lenti-HNF1A-shRNA ("HNF1AKD"). Error bars represent SEM. Two-tailed t-tests were used to generate *P*-values; **P*<0.01, ns= not statistically significant.

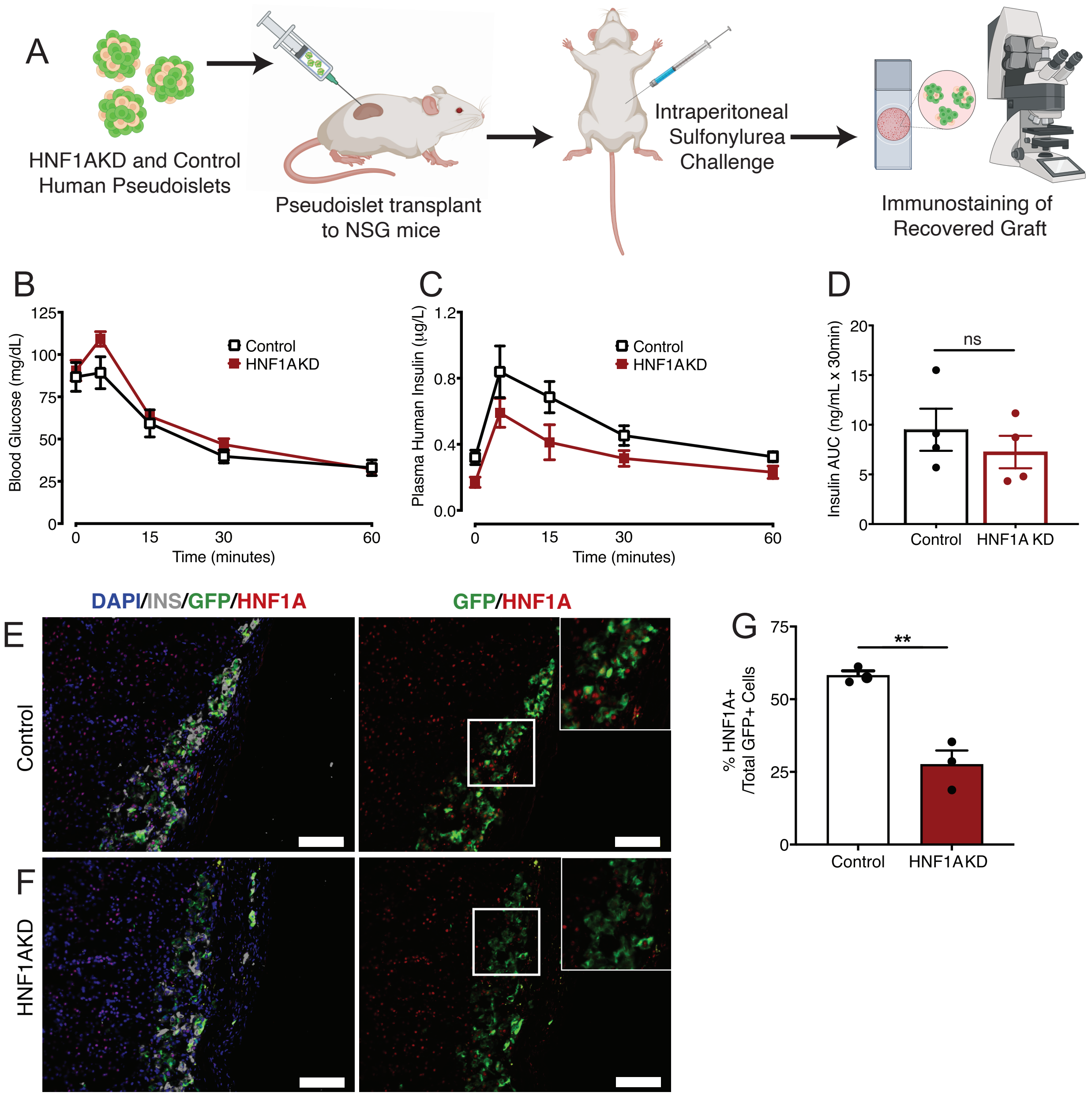


**Figure S2.** HNF1AKD is durable after transplantation and insulin phenotypes can be ameliorated with sulfonylurea treatment *in vivo*. (A) Schematic of experimental approach for pseudoislet transplantation to murine kidney capsules and characterization of long-term, *in vivo* phenotypes; "HNF1AKD" = pseudoislets transduced with lenti-HNF1A-shRNA; "Control" = pseudoislets transduced with lenti-Control-shRNA. (B) Blood glucose, (C) plasma human insulin levels, and (D) area under the curve of insulin excursion (not significant; $P$=0.44) upon intraperitoneal sulfonylurea challenge (n=4 mice, 3 human islet donors per condition). (E-G) Immunostaining of recovered grafts from mice transplanted with (E) Control or (F) HNF1AKD pseudoislets; scale bar= 100µm. (G) Quantification of $GFP^{+}HNF1A^{+}$ cells from grafts. Data are presented as mean values ± SEM. Two-tailed t-tests were used to generate $P$-values; **$P$<0.01.

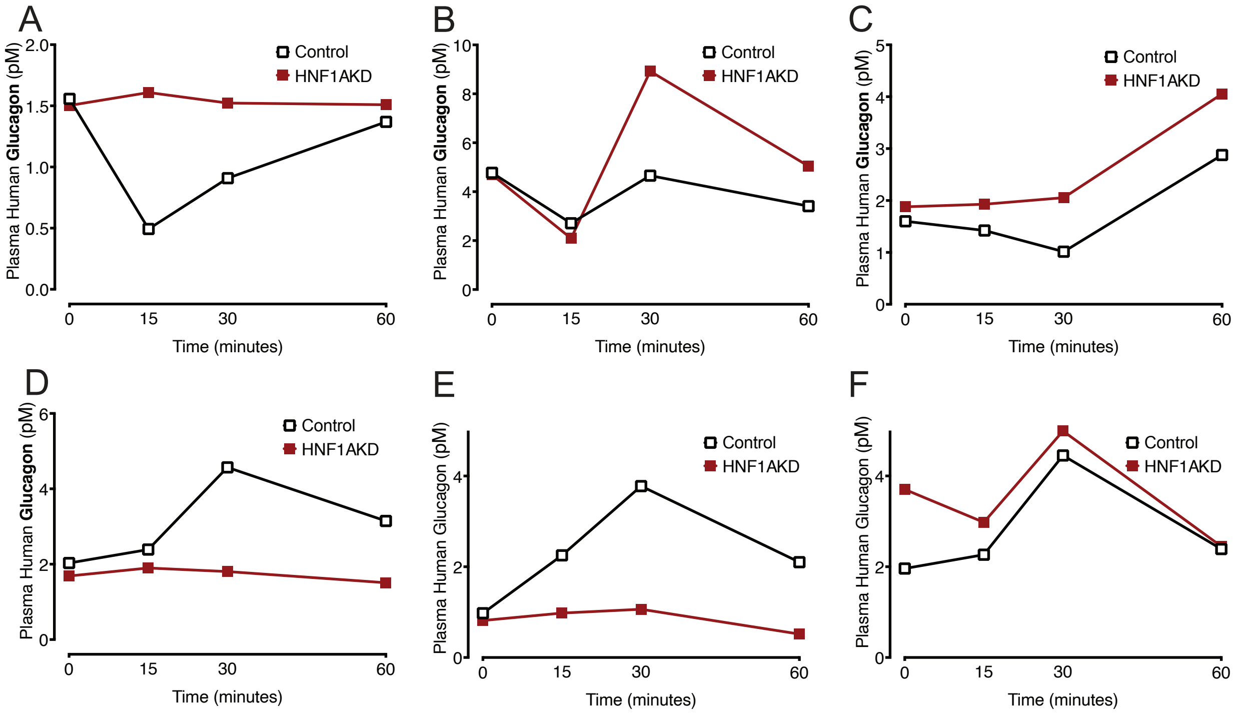


**Figure S3.** *HNF1A* suppression leads to dysregulated glucagon secretion after one month *in vivo*. Plasma glucagon excursion curves upon intraperitoneal (A-C) glucose or (D-F) insulin challenge after transplantation to GKO NSG mice. The graphs are data from 3 distinct human donors transplanted to 3 pairs of NSG-GKO mice (each mouse received either control or HNF1AKD pseudoislets from the same human donor): (A, D) donor 1, (B, E) donor 2, and (C, F) donor 3.

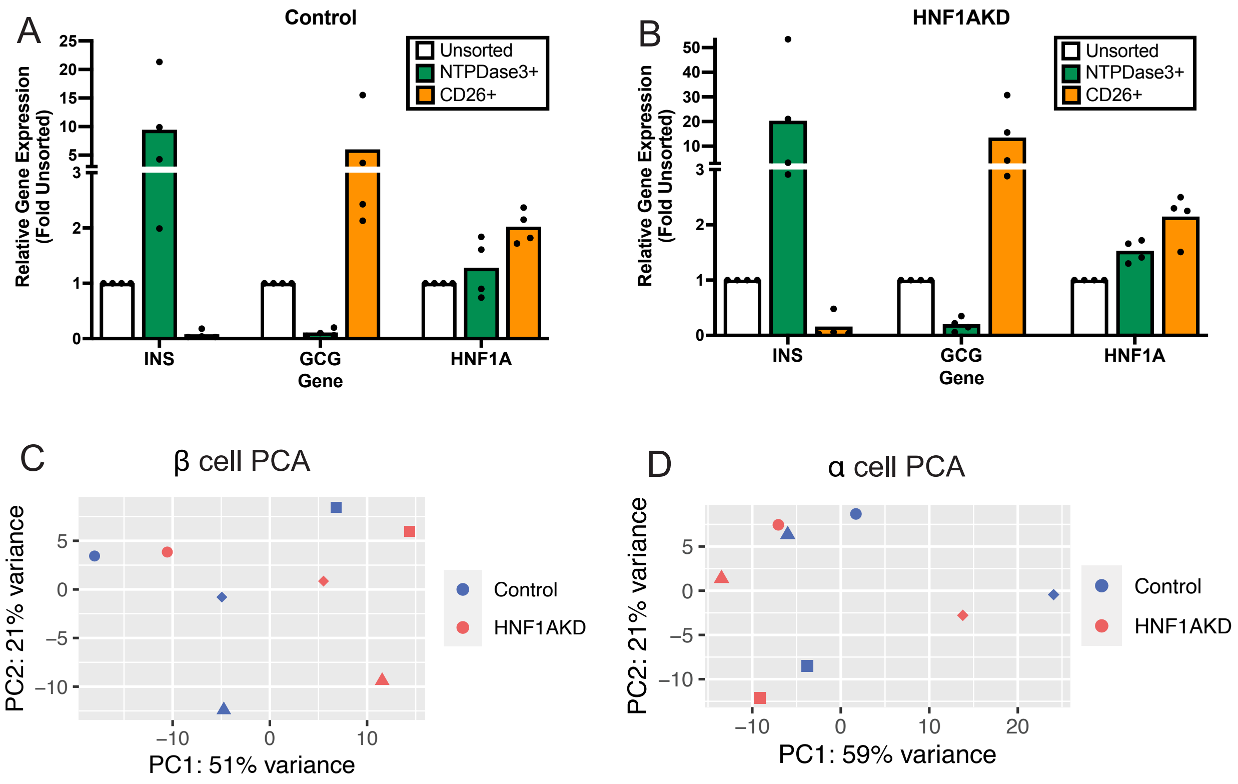


**Figure S4.** Fluorescence-activated cell sorting (FACS) of pseudoislets enriches β and α cell populations for RNA sequencing analysis. (A-B) RT-qPCR demonstrated enrichment of insulin (*INS*) with depletion of glucagon (*GCG*) in GFP$^+$HPi2$^+$NTPDase3$^+$ versus unsorted fractions and enrichment of glucagon with depletion of insulin in GFP$^+$HPi2$^+$CD26$^+$ fractions from sorted (A) Control and (B) HNF1AKD pseudoislet samples (n=4 distinct donors). (C-D) Principal Component Analysis (PCA) of RNA-sequencing results from sorted (C) β (NTPDase3$^+$CD26$^-$) and (D) α (CD26$^+$NTPDase3$^-$) cells of Control (blue) and HNF1AKD (salmon) pseudoislet samples (n=4 donors; distinct donors are indicated by different shapes).

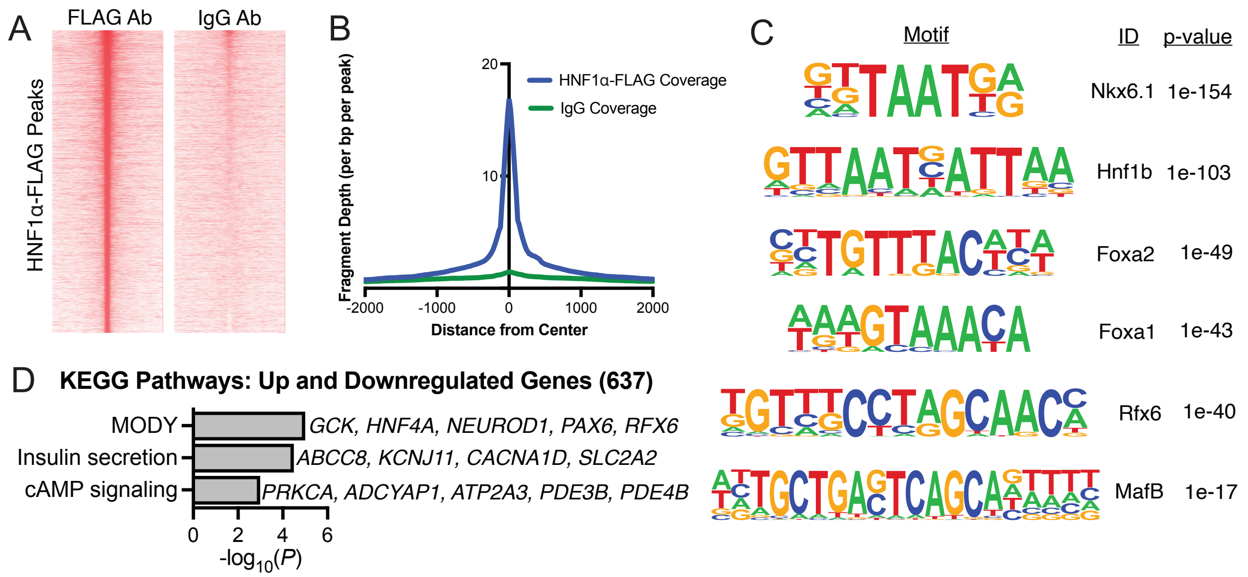


**Figure S5.** CUT&RUN identifies direct binding targets of HNF1α in primary human islet cells. (A) Heatmap visualization of CUT&RUN peaks and (B) histogram plots of averaged reads demonstrated enrichment of read densities in HNF1α-FLAG CUT&RUN library peak centers versus minimal enrichment at these sites for IgG control samples. (C) HOMER motif analysis identified that HNF1α-FLAG-bound genomic peaks were significantly enriched for pancreatic transcription factor motifs (NKX6.1, HNF1b, FOXA2, FOXA1, RFX6 and MAFB) previously shown to be enriched alongside HNF1α in pancreatic islet enhancer clusters. (D) KEGG pathways enriched in putative HNF1α target genes, which were identified by the intersection of HNF1α-FLAG CUT&RUN and HNF1AKD RNA-seq gene sets.
